# The RNA editing enzyme ADARB1 is readily detectable in primary auditory neurons and provides a means for automated counting

**DOI:** 10.64898/2026.03.26.714550

**Authors:** Garner C. Fincher, Punam Thapa, Steven C. Gressett, Bradley J. Walters

## Abstract

Spiral ganglion neurons (SGNs) are the primary auditory afferents in the inner ear. These neurons degenerate in response to a number of conditions, including auditory neuropathies, concussions, and aging. Research to assess the extent of degeneration and to test the efficacy of protective or rehabilitative strategies requires quantification of SGNs from tissue sections. However, manual counting of SGNs can be arduous and time-consuming due to dense crowding and the lack of reliable nuclear-specific labels. SGNs receive afferent input via GluA2-containing AMPA receptors. As the *Gria2* transcripts that code for GluA2 must undergo RNA editing to ensure calcium impermeability, we hypothesized that SGNs would express high levels of the adenosine deaminase acting on RNA (ADAR) enzyme ADARB1. Here we confirm enriched expression of *Adarb1* in SGNs via *in situ* hybridization and show that anti-ADARB1 antibodies robustly label the nuclei of both type I and type II SGNs in cochlear sections from young and aged mice. Neuronal specificity was confirmed using antibodies against neurofilament heavy chain (NFH), human antigen D (HuD), GATA binding protein 3 (GATA3), and SRY-box 2 (SOX2). A blinded investigator manually counted SGNs via NFH staining, and these were compared to automated counts of ADARB1-positive nuclei using the analyze particles function in ImageJ. A concordance correlation coefficient and Bland-Altman analysis demonstrated strong agreement between the manual and automated counts. Additionally, immunolabeling of ADARB1 in macaque and human temporal bone sections confirm robust labeling of SGN nuclei, suggesting broad utility of ADARB1 immunolabeling for automated counts of SGNs across species.

## INTRODUCTION

In contemporary auditory research, the counting of spiral ganglion neurons (SGNs) is essential. SGNs are the primary neurons that transmit auditory signals from the sensory hair cells of the cochlea to the brainstem. Moreover, they are often affected in sensorineural hearing loss (SNHL) which is a widespread phenomenon and a major human health issue. While sensory hair cell loss is viewed as the primary pathology for acquired hearing loss, degeneration of the SGNs has also been observed in a variety of contexts. Among the more prominent, recent examples are reports that suggest SGN loss may be the primary auditory pathology related to mild traumatic brain injuries or concussions (Ishai et al., 2018; Penn et al., 2023). SGN loss has also been a well-documented feature of aging (Bao & Ohlemiller, 2010; Perez et al., 2011). In addition, there are several auditory neuropathies that affect SGNs, including auditory neuropathy spectrum disorder, cochlear nerve deficiency, and congenital and idiopathic cases where degeneration of SGNs leads to reduced hearing or sound processing (Rapin & Gravel, 2003; Roche et al., 2010; Sagers et al., 2017; Starr et al., 2003). In many of these cases functional loss occurs despite the persistence of cochlear hair cells However, in cases of SNHL where sensory hair cell loss is the primary injury, SGN loss also occurs either acutely or secondarily. Thus, noise exposures, genetic factors, ototoxic medications such as aminoglycosides and platinum-based chemotherapeutics, inflammation and viral infection, and metabolic disturbances are all known causes of hair cell loss that can also lead to SGN loss(Alam et al., 2007; G Jansen et al., 2013; Huth et al., 2011; Jiang et al., 2017; Leake & Hradek, 1988; Tan & Vlajkovic, 2023). As a result, there has been increasing interest in quantifying the survival of SGNs in response to these many and varied environmental and genetic factors, and correspondingly increased interest in quantifying SGN numbers in response to interventions aimed at preventing or reversing cell loss. Historically, the counting of SGNs has been a time-consuming process, needing to be done manually, due to the relatively crowded nature of SGNs in Rosenthal’s canal and the lack of established nucleus-specific markers that would otherwise present as discreet or spatially separated.

In other parts of the nervous system, labeling with anti-NeuN antibodies has often been used for neuronal quantification. NeuN was the abbreviation given to an antigen to which antibodies were raised that seemed to selectively label Neuronal Nuclei (NeuN) throughout most of the brain (Mullen et al., 1992; Wolf et al., 1996). The protein bound by these antibodies has since been identified as RBFOX3, an RNA binding protein that does appear to be enriched in neurons as compared to other cells in many parts of the central nervous system (Kim et al., 2009; Wolf et al., 1996). However, NeuN immunolabeling has not been widely adopted for SGNs likely due to difficulties with NeuN labeling therein. Instead, over the past several decades, other neuronal markers have become established in the auditory field such as neurofilament heavy chain (NF-H), beta-III tubulin (Tuj1), and human antigen D (HuD) which is an established neuronal marker now known to be ELAVL4 (Berglund & Ryugo, 1991; Brooks et al., 2020; Coate & Kelley, 2013; Jyothi et al., 2010; Nayagam et al., 2011; Vanevski & Xu, 2015). Indeed, our personal experience, and communications with colleagues, suggest that previous generations of NeuN antibodies did not adequately label SGNs, while more recently developed NeuN antibodies that do label SGNs do not appear to be restricted to cell nuclei ((Elliott et al., 2021; Filova et al., 2020, 2022) and also see Figure 1). This could be for any of several reasons, including: possibly lower levels of RBFOX3 in SGN nuclei as compared to other neuron types, possible cytoplasmic localization and function of RBFOX3 in SGNs, and/or potentially broader or less-specific labeling of the newer antibodies. In any case, these factors suggest NeuN immunolabeling may still offer limited utility for SGN quantification. Similarly, other neuronal markers adopted by the field appear limited for this purpose in that they label the somal cytoplasm (e.g. HuD) and/or neuronal processes (e.g., Tuj1, NFH)(Berglund & Ryugo, 1991; Brooks et al., 2020; Coate & Kelley, 2013; Vanevski & Xu, 2015). This non-nuclear staining leads to crowded fields of view and difficulty in delineating individual neurons, particularly in automated workflows, which prompted us to identify a marker specific for the SGN cell nuclei.

**Figure 1:**
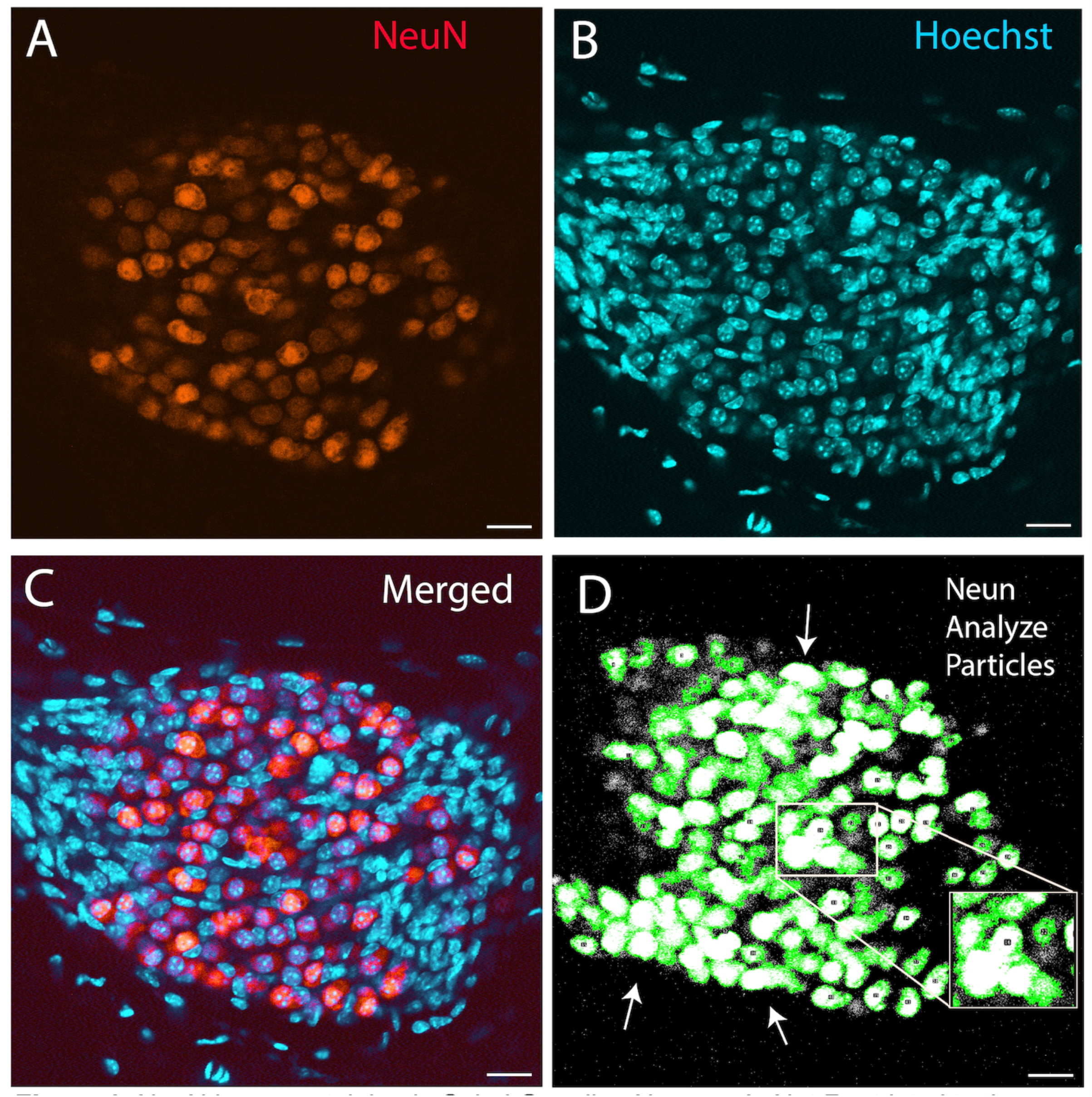
NeuN immunostaining in spiral ganglion neurons is not restricted to the nucleus. (A-C) Representative confocal images of spiral ganglion neurons (SGNs) from P21 mouse cochleae immunostained for (A) NeuN (orange), an established marker of neurons expected to be specifically localized in cell nuclei, and (B) Hoechst stain (cyan), an established marker of cell nuclei. C) Co-localization shows NeuN immunolabeling is often present within the Hoechst-positive nuclear region, but also visibly extends into the surrounding cytoplasm suggesting that, in SGNs, NeuN is not exclusively nuclear. D) NeuN labeled images like the one in (A) were made binary in ImageJ and the analyze particles function was used to quantify the numbers of SGNs. As noted (white arrows, box) the extranuclear nature of the NeuN staining led to many “fused” cells and undercounting. Scale bar = 20 µm

It is well-established that A-to-I editing of the *Gria2* transcript that codes for the GluA2 receptor subunit is required in order for GluA2-containing AMPA receptors to be impermeable to calcium (Konen et al., 2020; Seeburg et al., 1998; Sommer, Kohler, et al., 1991). This editing is accomplished by the ADARB1 enzyme (a.k.a. ADAR2) wherein its deletion in mice leads to seizures and perinatal lethality due to the failure of such AMPA receptors to become calcium impermeable (Higuchi et al., 2000; T. Y. Tan et al., 2020). As SGNs are glutamatergic, and readily express the GluA2 subunit, we hypothesized that SGNs would express high levels of ADARB1. Indeed, RNA-seq datasets (Li et al., 2020; Shrestha et al., 2018) suggest that *Adarb1* is prevalent in murine SGNs, with a notably high neuron:glia expression ratio. Further, ADARB1 protein is expected to be enriched specifically in the nuclei of the SGNs as the nucleus is the primary site of RNA editing by ADARs, and similar nuclear localization of ADARB1 is typically seen in neurons in the CNS (Behm et al., 2017; Moore et al., 2019).

To determine whether ADARB1 is localized in SGN nuclei and whether this property could be used to automate SGN quantification, we collected temporal bone sections from mice at neonatal, young adult, and advanced ages and performed immunofluorescent labeling with ADARB1 antibodies. Co-labeling with antibodies against NFH and HuD allowed for confirmation of neuronal identity and specificity, and co-labeling with a GATA3 antibody further confirmed ADARB1 presence in type II as well as type I SGNs. Additional co-labeling experiments with anti-SOX2 revealed no strong ADARB1 signal in glial cells, thus suggesting that ADARB1 immunofluorescent signal could be used to identify SGN neurons specifically. We then compared manual counts of neurons by an investigator who used NFH labeling to those of an investigator who used ADARB1 labeling and the analyze particles function in ImageJ. Similar numbers of neurons were counted by both methods. Finally, archived temporal bone sections from macaques and human donors were immunolabeled for ADARB1 and similarly demonstrated strong nuclear labeling in the SGNs suggesting ADARB1 labeling as a promising method for SGN quantification across taxa.

## Materials and Methods

### 1. Animal Care and Housing

Experiments were conducted on CD-1 mice at postnatal day 0 (P0) and P21, and on C57Bl/6 mice aged to 24 months. All mice used for this study were housed in the animal care facilities of the University of Mississippi Medical Center (UMMC) in a temperature-controlled environment with *ad libitum* food and water access and on a 12:12 light: dark cycle. All experimental procedures in the present study were approved by the Institutional Animal Care and Use Committee (IACUC) at UMMC. For the detection of ADARB1 at P0 and co-localization of ADARB1 with neuronal and glial markers, we used sections from at least 4 different mice from multiple litters, both male and female. For comparisons of manual counting to automated counting in P21 mice, we used sections from at least three male and three female mice (n=6) again taken from multiple litters. Similarly, temporal bone sections from more than 4 mice, both males and females, were used for the 24-month-old C57Bl/6 samples.

### 2. Tissue Preparation

Temporal bones were resected and then fixed in 4% paraformaldehyde (PFA) for three hours at room temperature (RT). Following fixation, temporal bones from P21 mice and 24-month-old mice were decalcified in 0.25M EDTA at PH 7.5, with the P21 samples immersed for 24 hours at 4°C and the 24-month samples for 96 hours. Temporal bones were then stored in 1x PBS at 4°C until ready for sectioning. Cochleae were embedded in 5% agarose (BP1356-500, Fisher bioreagents) using tissue embedding molds (Peel-A-Way, Polysciences, inc.). Once solidified, gel blocks were affixed to the sample holder of a vibrating microtome (5100mz, Campden Instruments) with Gorilla Glue Super Glue Gel and sectioned at a thickness of 60 - 80 µm. Next, we incubated the sections in a blocking buffer consisting of 1% triton-X, 10% bovine serum albumin (fraction V), and 10% normal goat serum in 1x Tris buffered saline (TBS) for one hour at RT. Samples were then incubated in primary antibodies overnight at 4°C and then sections were washed with TBS. The sections were incubated with secondary antibodies for 2 hours at RT, washed with TBS then immersed in Hoechst (1:1000, Invitrogen H3570) for 30 minutes at RT. For samples immunolabeled with NFH, primary and secondary antibodies were applied at 37°C. For GATA binding protein 3 (GATA3) staining, the sections were pre-treated with high pH antigen unmasking solution (Vector laboratories, H-3301, Lot ZF0613) for 20 minutes at 85°C in a humidified chamber and washed with TBS. Sections were incubated in Image-iT FX signal enhancer (Invitrogen I36933) for 30 minutes at RT and then washed with TBS prior to staining. Human and macaque sections were obtained as described previously (Ghosh, Wineski, et al., 2022). Briefly, archived sections (> 5 years in storage) were retrieved from −80°C and were washed and equilibrated with ultrapure water at RT to remove optimum cutting temperature compound (OCT). Sections were bounded using a hydrophobic pen then washed in PBS and incubated in antibodies with intervening washes as described above. For the immuno-histochemistry technique, the primate samples were washed in PBS then incubated for 2 hours at RT in blocking buffer containing 5% triton-X, then followed by 1% triton-X, 1% BSA incubation for an hour and 10% normal serum followed by 72 hours incubation at 4°C in primary antibodies to yield better results. Of note, some of the human sections were probed twice or were probed for longer periods of time with higher concentration of antibody due to high background, autofluorescence, and/or poorer performance of the antibody labeling. In the case of re-probing, samples mounted in Fluorogel/PBS/Glycerol had their coverslips removed and the sections re-run through primary and secondary antibody steps. Exacerbated steps for some human sections thus included 3 days of incubation in primary antibody at RT, and ADARB1 antibody concentrations as high as 1:50. All primary and secondary antibodies used are listed in Table 1. RNAscope samples were obtained in the manner described above: P21 temporal bones were decalcified in 0.25 M EDTA at PH 7.5 for 24 hours at 4°C and subsequently embedded in 5% agarose with eventual subsequent vibratome sectioning as described above. Tissue sections were then processed for RNAscope experiments conducted as previously described (Ghosh, Casey, et al., 2022). The RNAscope probes used for these experiments are outlined in table 2.

**Table 1.**
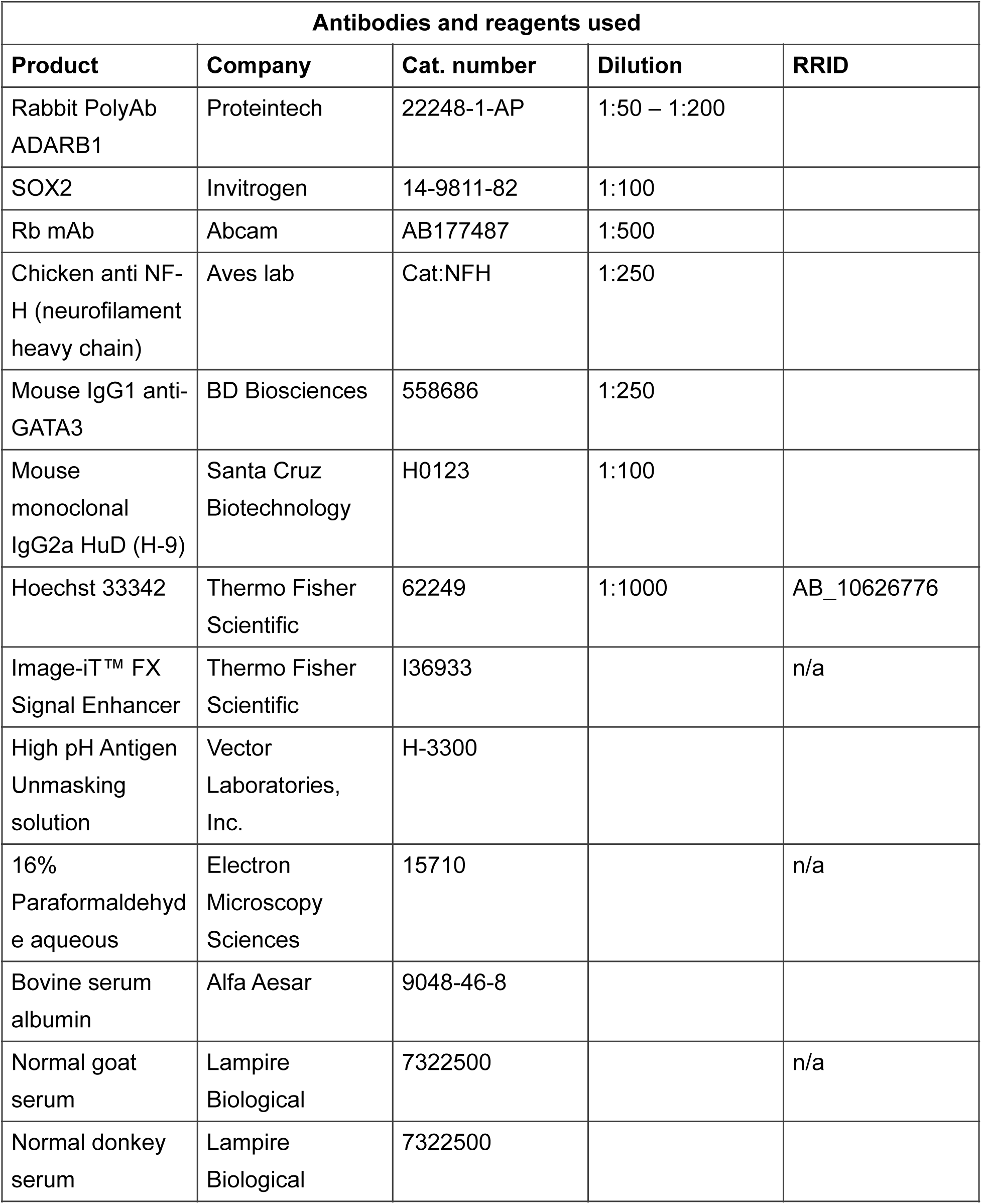
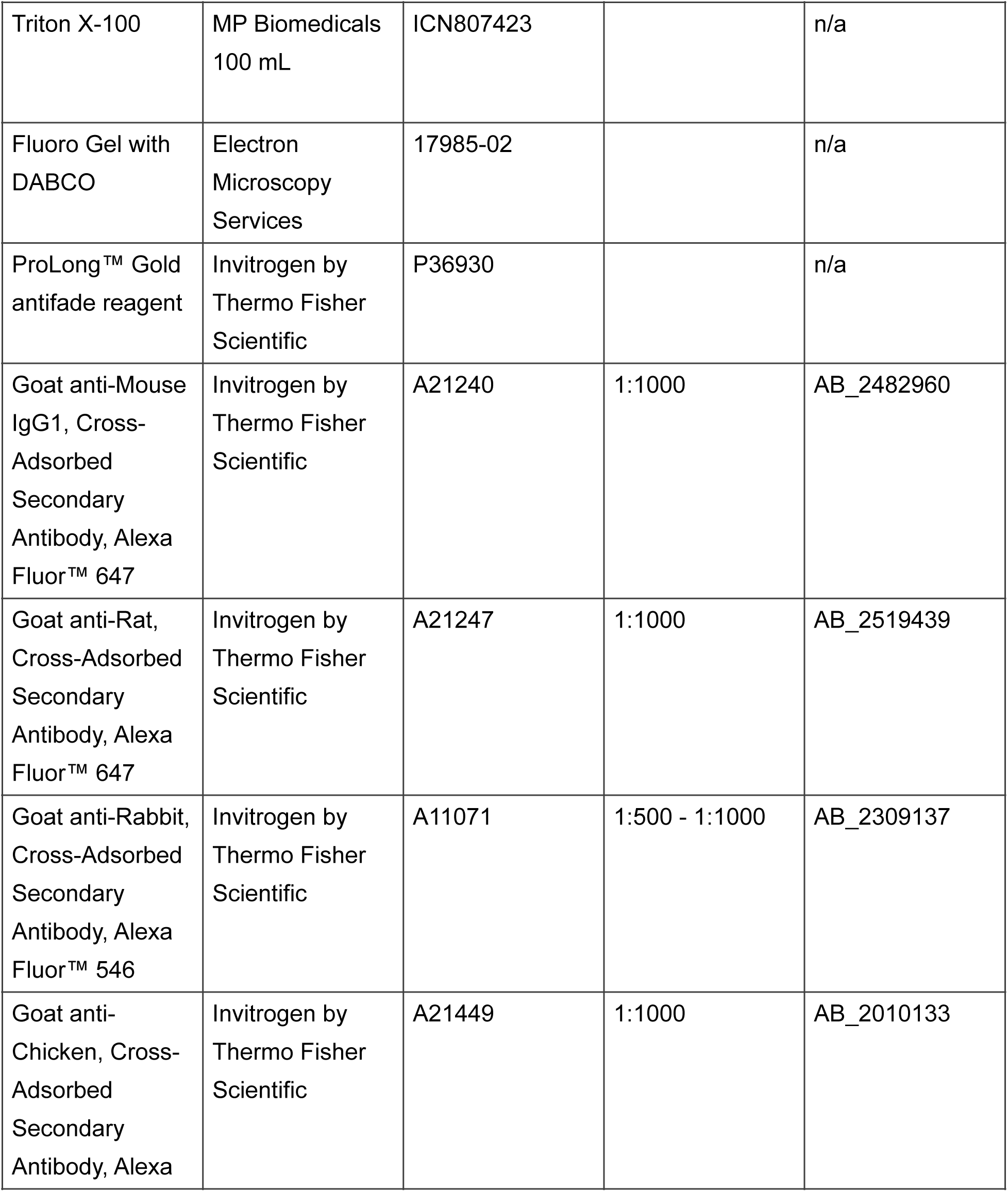
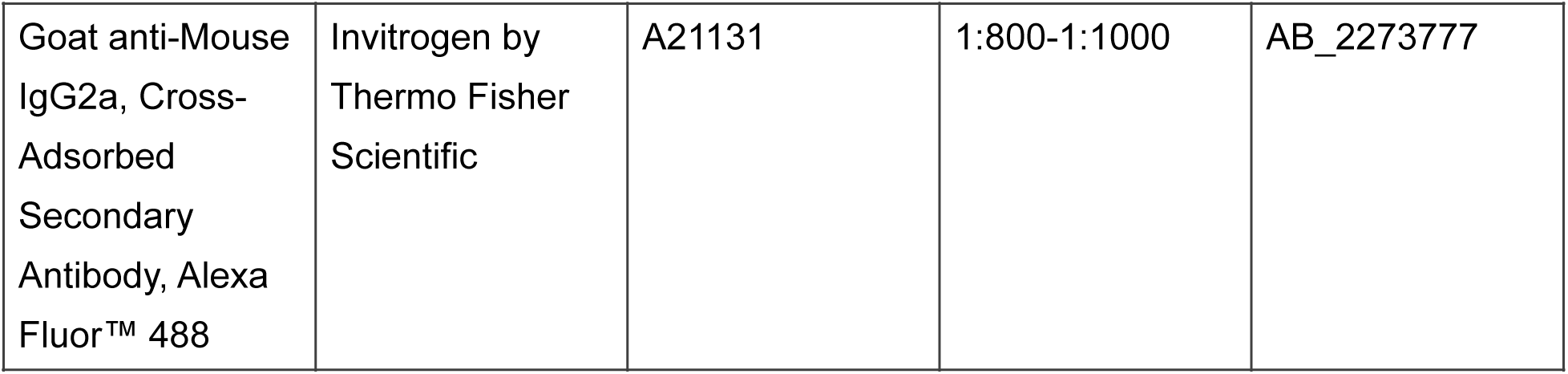

| **Antibodies and reagents used** |  |  |  |  |
| --- | --- | --- | --- | --- |
| **Product** | **Company** | **Cat. number** | **Dilution** | **RRID** |
| Rabbit PolyAb ADARB1 | Proteintech | 22248-1-AP | 1:50 – 1:200 |  |
| SOX2 | Invitrogen | 14-9811-82 | 1:100 |  |
| Rb mAb | Abcam | AB177487 | 1:500 |  |
| Chicken anti NF-H (neurofilament heavy chain) | Aves lab | Cat:NFH | 1:250 |  |
| Mouse IgG1 anti-GATA3 | BD Biosciences | 558686 | 1:250 |  |
| Mouse monoclonal IgG2a HuD (H-9) | Santa Cruz Biotechnology | H0123 | 1:100 |  |
| Hoechst 33342 | Thermo Fisher Scientific | 62249 | 1:1000 | AB_10626776 |
| Image-iT™ FX Signal Enhancer | Thermo Fisher Scientific | I36933 |  | n/a |
| High pH Antigen Unmasking solution | Vector Laboratories, Inc. | H-3300 |  |  |
| 16% Paraformaldehyde aqueous | Electron Microscopy Sciences | 15710 |  | n/a |
| Bovine serum albumin | Alfa Aesar | 9048-46-8 |  |  |
| Normal goat serum | Lampire Biological | 7322500 |  | n/a |
| Normal donkey serum | Lampire Biological | 7322500 |  |  |
| Triton X-100 | MP Biomedicals<br>100 mL | ICN807423 |  | n/a |
| Fluoro Gel with DABCO | Electron Microscopy Services | 17985-02 |  | n/a |
| ProLong™ Gold antifade reagent | Invitrogen by Thermo Fisher Scientific | P36930 |  | n/a |
| Goat anti-Mouse IgG1, Cross-Adsorbed Secondary Antibody, Alexa Fluor™ 647 | Invitrogen by Thermo Fisher Scientific | A21240 | 1:1000 | AB_2482960 |
| Goat anti-Rat, Cross-Adsorbed Secondary Antibody, Alexa Fluor™ 647 | Invitrogen by Thermo Fisher Scientific | A21247 | 1:1000 | AB_2519439 |
| Goat anti-Rabbit, Cross-Adsorbed Secondary Antibody, Alexa Fluor™ 546 | Invitrogen by Thermo Fisher Scientific | A11071 | 1:500 - 1:1000 | AB_2309137 |
| Goat anti-Chicken, Cross-Adsorbed Secondary Antibody, Alexa | Invitrogen by Thermo Fisher Scientific | A21449 | 1:1000 | AB_2010133 |
| Goat anti-Mouse IgG2a, Cross-Adsorbed Secondary Antibody, Alexa Fluor™ 488 | Invitrogen by Thermo Fisher Scientific | A21131 | 1:800-1:1000 | AB_2273777 |

**Table 2.**

| RNA Scope In Situ Hybridization Probes |  |  |  |  |
| --- | --- | --- | --- | --- |
| Probes | Species | Reference no. | Lot no. | Supplier |
| *mm-Adarb1* | *Mus. Musc.* | 517431 | 221538 | ACD-Bio |
| *mm-negative control-DAPB* | *Mus. Musc* | 310043 | 2013100 | ACD-Bio |

### 3. Confocal Microscopy and Image Processing

After immunolabeling and/or RNAscope, sections were washed with TBS and then coverslipped using Prolong gold antifade reagent (Life Technologies Corporation, P36930, Lot 2897809) or using fluorogel with DABCO (Electron Microscopy Sciences, 17985) and 50% glycerol in PBS at a 1:1 ratio. Slides were kept for 2-3 days in the dark to allow mounting media to set, after which samples were imaged using a Zeiss LSM880 confocal microscope with a 20X dry or 40x oil objective. Image stacks (z-interval = 1 µm) were acquired of Rosenthal’s canal in the apex, middle, and basal turn of each temporal bone from mid-modiolar sections. Images used for counting were initially processed through ImageJ (https://imagej.net/software/fiji/downloads) where 2 dimensional images were created by maximum intensity projection (MIP) of optimal (6 or less) z-slices from each acquisition file. The channels were then separated using the split channels function so the different channel images would have the same ROI. The NFH channel images were then counted by a blinded investigator. The ADARB1 channel images were converted to binary data in ImageJ by using selecting: process > binary > make binary. The analyze particles function was then used to produce automated counts. Typically, a minimum pixel size was not required, however in some instances where signal bled-through, or background was prominent, we set a threshold to exclude small artifacts. This was generally solved by setting the minimum Size for the particle analyzer to 9-14µm^2^ and leaving the maximum set to infinity.

### 4. Statistical Analysis

Inter-rater reliability between the manual and automated counts was assessed using each analyzed image as a data point. A Bland-Altman Plot and Lin’s concordance correlation coefficient were calculated in R using the SimplyAgree package(Caldwell, 2022).

## Results

### 1. Anti-NeuN labels the nuclei and soma of SGNs in the mouse inner ear

The counting of SGNs has historically been both a time- and labor- intensive process due to the crowding observed with traditional markers such as NFH, Tuj1, and HuD which most prominently label the fibers and/or the soma. Anti-NeuN antibodies have been widely used in contemporary neuroscience research to label neuronal nuclei (Gusel’nikova & Korzhevskiy, 2015), however, not all neurons are labeled by NeuN antibodies, with Purkinje neurons, olfactory bulb mitral cells, and retinal photoreceptors (in mammals) being notable exceptions (Moon et al., n.d.; Mullen et al., 1992b; Saito et al., 2017). It is suspected that NeuN labeling of SGNs has been similarly limited in the past by either lower levels of RBFOX3 and/or poor sensitivity of previous generations of NeuN antibodies. Indeed, the first characterization of NeuN immunolabeling in the CNS was in 1992 (Mullen et al., 1992a), yet despite wide adoption in studies throughout the CNS in the 1990s, 2000s, and 2010s, there have been only a few reports demonstrating NeuN immunoreactivity in SGNs *in vivo*, and those have all come within the past few years(Elliott et al., 2021; Filova et al., 2020, 2022) Furthermore, during the intervening years, numerous reports were published that demonstrated useful immunoreactivities of antibodies against neurofilament proteins, calcium binding proteins, and other cytosolic, membrane bound, and synaptic proteins as markers for SGNs (Berglund & Ryugo, 1991; Coate & Kelley, 2013; Jyothi et al., 2010; Nayagam et al., 2011; Vanevski & Xu, 2015; Wolf et al., 1996), suggesting a strong desire to identify alternative markers. In the more recent publications that demonstrate NeuN immunoreactivity in the spiral ganglion (Elliott et al., 2021; Filova et al., 2020, 2022) the figures tend to be at lower power yet still suggest that NeuN labeling may not be nuclear-specific in primary auditory neurons, but likely also label the soma. This would be consistent with other reports that suggest NeuN may be present in the soma, or even more prominent in the cytosol rather than in the nuclei, of some neurons (Anderson et al., 2021). Thus, to determine the specificity of NeuN immunolabeling in the auditory periphery, we conducted immunofluorescent analysis on mouse temporal bone sections, with higher power (400X total magnification) confocal imaging focusing on spiral ganglion neurons (SGNs). As illustrated in Figure 1, NeuN immunofluorescence was consistently bright within the SGNs, but it was not confined to the nuclear compartment. Instead, NeuN immunoreactivity extended into the perinuclear cytoplasm and soma, suggesting that RBFOX3 may also be present in extranuclear compartments or that the antibody cross-reacts with cytoplasmic isoforms or related proteins. Across multiple sections and biological replicates, we observed this pattern of nuclear and somal labeling. This broader distribution seems to lessen the utility of NeuN as a strictly nuclear marker, posing challenges for automated quantification, as it hinders the ability to reliably distinguish or segment individual neurons, particularly in densely packed regions (see Figure 1D).

### 2. ADARB1 expression becomes prominent in SGNs as the inner ear matures

As noted, SGNs express the GluA2 receptor subunit at high levels (Matsubara et al., 1996; Ruel et al., 1999; Rutherford & Moser, 2016; Safieddine et al., 2012), and GluA2 (*Gria2)* transcripts must be edited by ADARB1 in order for the receptors to function properly (Burnashev et al., 1992; Higuchi et al., 1993; Sommer, Köhler, et al., 1991). Furthermore, RNA seq data from mice has shown that *Adarb1* expression increases dramatically during embryonic development of SGNs (Li et al., 2020) which is consistent with other reports demonstrating increasing ADARB1 immunoreactivity in neurons as they mature (Cuddleston et al., 2022; Dailamy et al., 2024). We therefore hypothesized that *in situ* hybridization and immunolabeling would reveal high levels of the RNA editing enzyme ADARB1 in mature SGNs. Thus, we sought to initially investigate *Adarb1* transcript expression and ADARB1 immunoreactivity at P21, a time when mice are first able to hear across a broad frequency range and the SGNs are known to be functional (Souchal et al., 2018). Our results suggest that *Adarb1* transcripts are indeed enriched in SGNs (Figure 2), and that ADARB1 immunolabeling is enriched specifically within SGN nuclei in CD1 mice at this age (Figure 3). We next sought to determine if ADARB1 immunoreactivity might be similarly robust in SGNs at an early postnatal age, prior to hearing onset, and thus investigated patterns of immunostaining at P0. At P0, ADARB1 expression is already fairly enriched in SGNs, but unlike at P21, it demonstrates a broader pattern across other cell types around Rosenthal’s canal (Figure 4), presumably mesenchymal cells (Rose et al., 2023) as well as cells in other parts of the sections including the cochlear lateral wall (not shown). This more widespread distribution suggests that, although neuronal expression is prominent, ADARB1 lacks strict cell-type specificity at this perinatal stage. By P21, ADARB1 expression persists at high levels in SGNs, but its detectability in non-neuronal cell types diminishes markedly relative to the SGNs (Figure 5). These patterns, along with the cited RNAseq data suggest that *Adarb1* is likely fairly ubiquitously expressed at baseline levels across cell types of the inner ear, but increasing levels in the SGNs likely eclipses detection in other cell types by P21. To confirm the specificity of ADARB1 immunolabeling in SGNs, we further characterized samples from P21 mice using antibodies with known neuronal or glial preferences in labeling. Indeed, the pattern of ADARB1 immunofluorescence is much more restricted to SGNs specifically at P21 (Figure 3). To determine whether ADARB1 persists in the cells surrounding Rosenthal’s canal, we increased laser power, detector gain, and image brightness and contrast so as to oversaturate the signal in the SGNs. This suggested that ADARB1 is not completely lost from other cell types, as some staining is detectable in cell nuclei outside of Rosenthal’s canal under these conditions (Figure 6). However, as illustrated, the signal-to-noise ratio of ADARB1 expression in SGN nuclei as compared to other cell types is remarkably high, and even when oversaturated seems to be absent from glial cells in Rosenthal’s canal thus allowing for SGN quantification without much interference.

**Figure 2:**
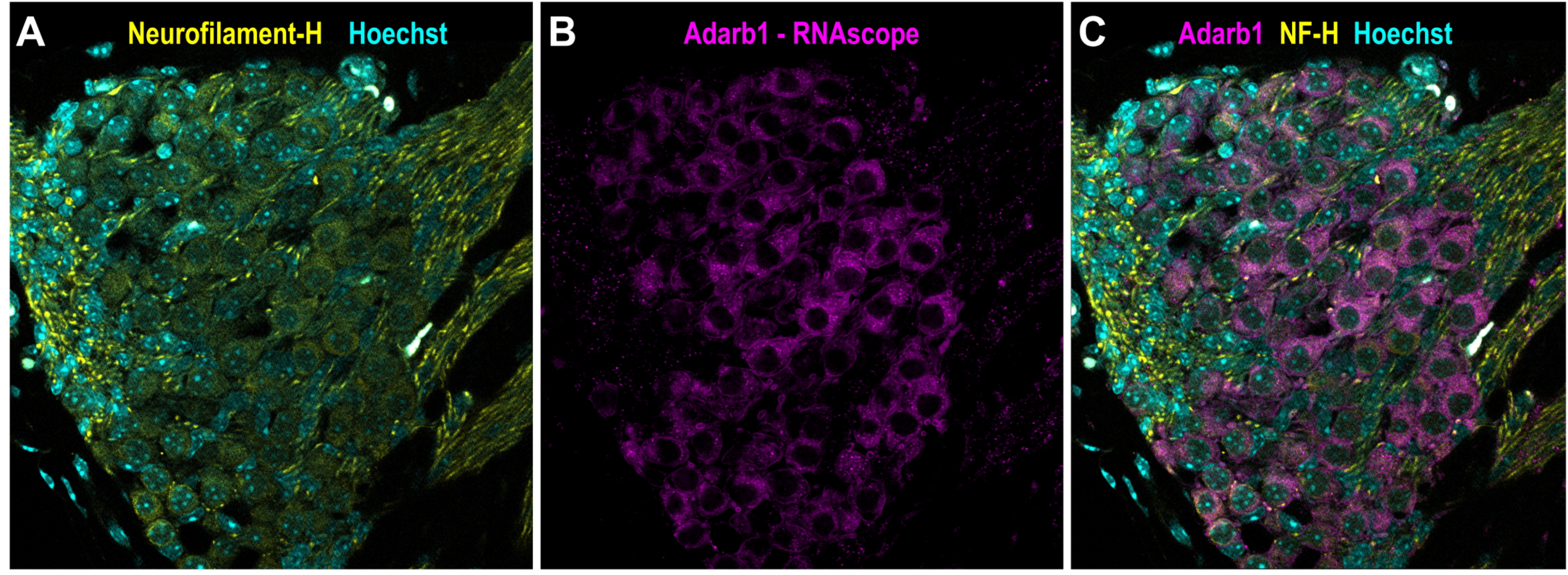
*Adarb1* (magenta) RNAscope *in situ* hybridization of temporal bone sections from mice at P21 suggests high levels of Adarb1 expression in SGNs within Rosenthal’s canal. (A-C) Representative confocal micrographs of the spiral ganglion in cross-section from a postnatal day 21 (P21) mouse cochlea immunolabeled with NFH (yellow) and Hoechst (cyan) suggest high levels of *Adarb1* expression (magenta) in NFH-positive SGNs. Scale bar = 20 µm

**Figure 3:**
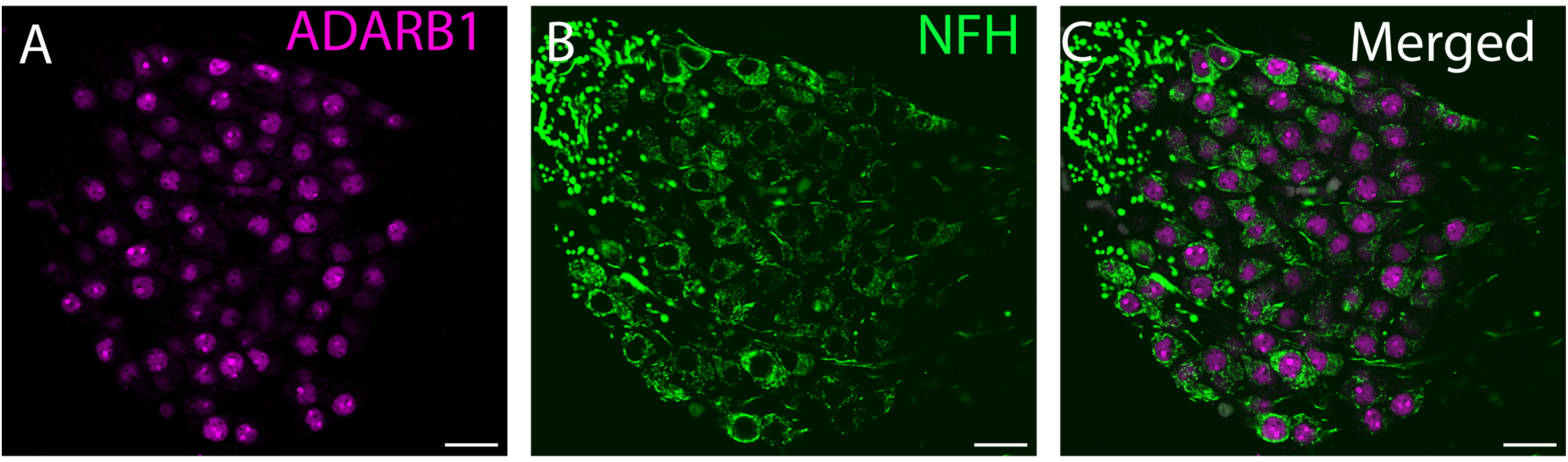
ADARB1 immunostaining consistently labels the nuclei of all neurofilament heavy chain (NFH)-positive spiral ganglion neurons (SGNs), underscoring its reliability for automated quantification. (A–C) Confocal micrographs of Rosenthal’s canal in a section from a postnatal day 21 (P21) mouse cochlea immunolabeled with ADARB1 (magenta) and NFH (green). (A) ADARB1 displays robust nuclear localization within SGNs, providing clear delineation of individual neuronal nuclei throughout the section. (B) NFH, an established neuronal marker routinely used in manual SGN quantification, exhibits strong labeling of the SGN fibers and soma. (C) The merged image reveals complete overlap between ADARB1 and NFH in the same cells, indicating that all NFH-positive neurons also exhibit ADARB1 nuclear staining. Scale bar = 20 µm

**Figure 4:**
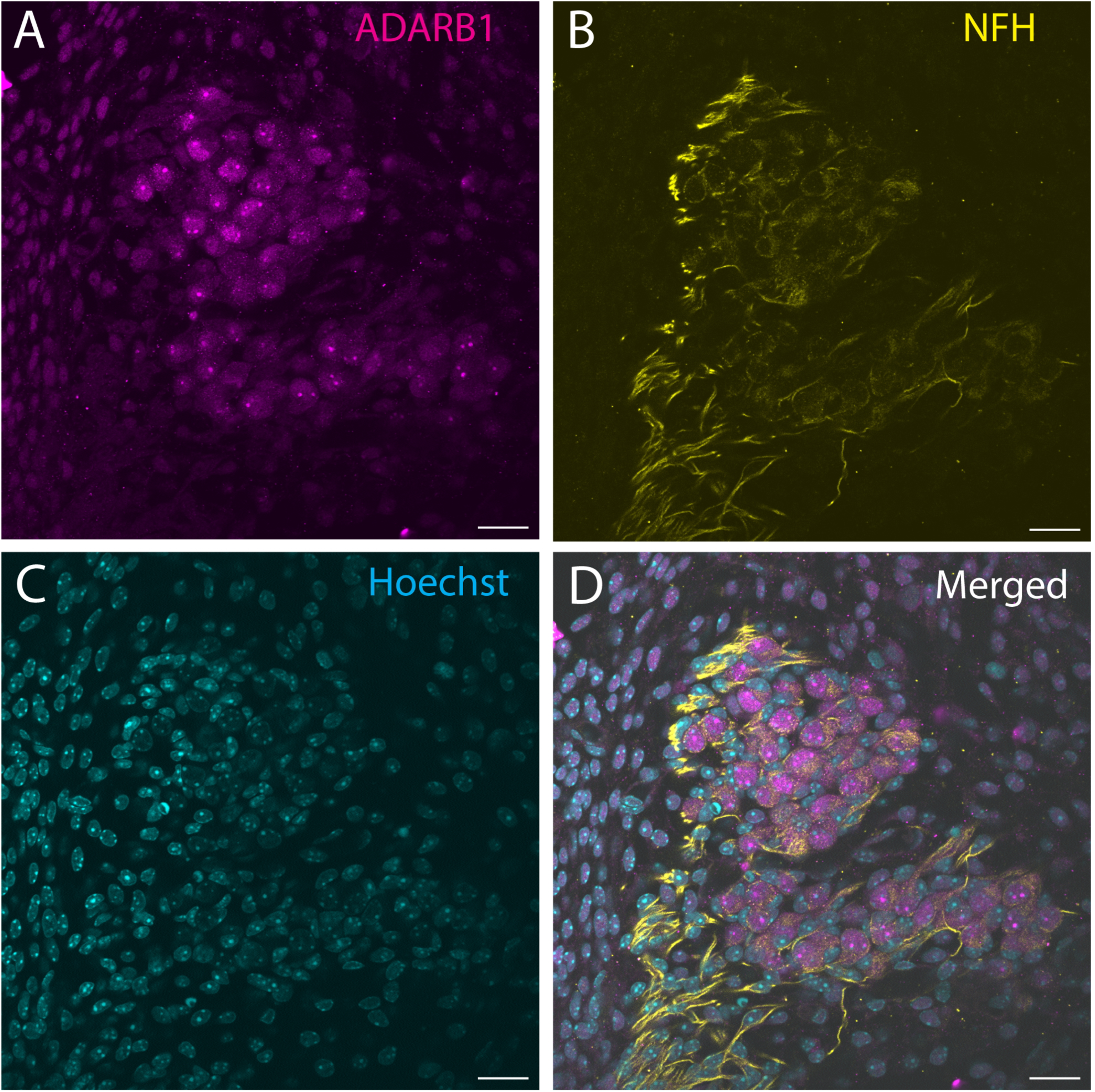
ADARB1 immunolabeling of temporal bone sections from mice at P0 suggests moderate ADARB1 expression in SGNs and less intense immunofluorescence in other cell types outside of Rosenthal’s canal. (A-C) Confocal micrographs of a P0 mouse temporal bone section immunolabeled for ADARB1 (magenta) and co-labeled with Hoechst (blue) indicate nuclear localized ADARB1 in SGNs and also in cells surrounding Rosenthal’s canal compared to P21 mice, where ADARB1 is so enriched in SGNs that its immunofluorescence is largely undetectable in other cell types. Scale bar = 20 µm

**Figure 5:**
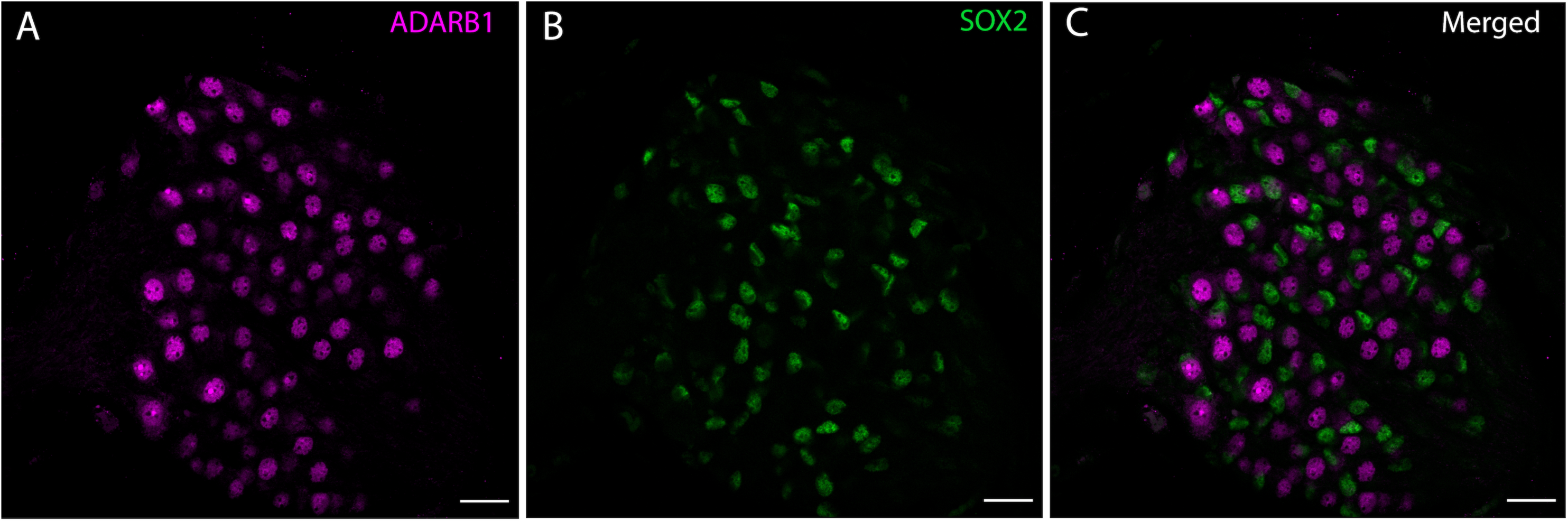
Immunostaining for ADARB1 and SOX2 in P21 mouse temporal bone sections demonstrates that ADARB1 immunofluorescence is not prominent in glial nuclei within Rosenthal’s canal. (A–C) Representative confocal micrographs of the spiral ganglion in cross-section from a postnatal day 21 (P21) mouse cochlea immunolabeled with ADARB1 (magenta) and SOX2 (green) suggest no strong colocalization between ADARB1-positive and SOX2-positive nuclei. Scale bar = 20 µm

**Figure 6:**
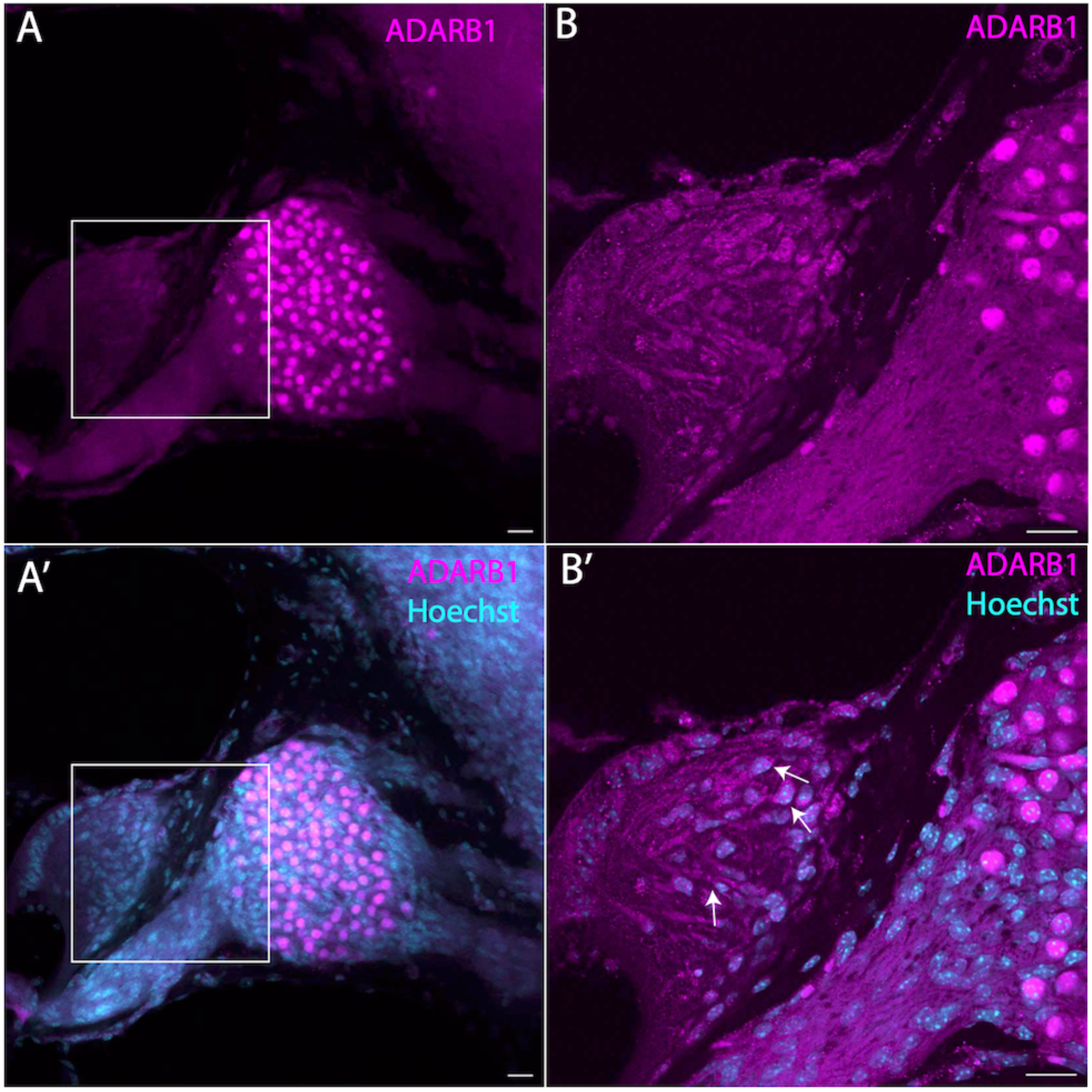
Increasing the intensity of ADARB1 immunofluorescence reveals detectable signal in cells outside of Rosenthal’s canal in P21 mice. (A–A’) Representative 20× images of Rosenthal’s canal show a high degree of specificity for ADARB1 expression in spiral ganglion neurons (SGNs). (B–B’) 40× images further highlight the specificity of ADARB1 in SGNs which generally must be oversaturated to reveal detectable ADARB1 nuclear-localized signal in cells located outside of Rosenthal’s canal. For example, colocalization of ADARB1 (magenta) and Hoechst (cyan)suggest a number of ADARB1-positive cells in the spiral limbus some of which are denoted by white arrows. These findings suggest that, although ADARB1 is highly specific to SGNs, low levels of expression are also present in other cell types. Scale bar = 20 µm

### 3. Co-labeling for SOX2, NFH, HuD, and GATA3 confirms that ADARB1 immunofluorescence allows for labeling of both type I and type II SGNs

A key objective of our study was to confirm the neuronal enrichment of ADARB1 and to determine if it is highly expressed in both type I and type II SGNs. To verify this, we first co-labeled sections with SOX2 which is known to be enriched in glial cells and not readily detectable in SGNs (Nishimura et al., 2017). These experiments revealed separate and distinct labeling of ADARB1 and SOX2 with no obvious colocalization, suggesting that ADARB1 is not highly expressed in the glial cells of Rosenthal’s canal at P21 (see Figure 7). In order to confirm the neuronal specificity of ADARB1 immunofluorescence, we examined its colocalization with the conventional neuronal markers HuD and NFH. Indeed, numerous reports have demonstrated these as markers for SGNs (Berglund & Ryugo, 1991; Brooks et al., 2020; Coate & Kelley, 2013; Vanevski & Xu, 2015). ADARB1 positive nuclei were consistently colocalized within NFH-positive signal (Figure 2) or within HuD positive soma (Figure 7), confirming their enriched presence in SGNs. As there are two primary types of SGNs, type I and type II, and as type II neurons comprise only a small portion of total SGNs, we also sought to confirm ADARB1 immunoreactivity in type II SGNs specifically. To do this, we co-labeled samples with antibodies against ADARB1 and GATA3, the latter being a transcription factor that has been shown to demarcate type II SGNs in the mature mouse inner ear (Nishimura et al., 2017). As illustrated in Figure 8, ADARB1 immunofluorescence was prominent in all GATA3 positive SGNs suggesting that ADARB1 immunolabeling is effective for the identification of all afferent SGNs regardless of whether they are type I or type II.

**Figure 7:**
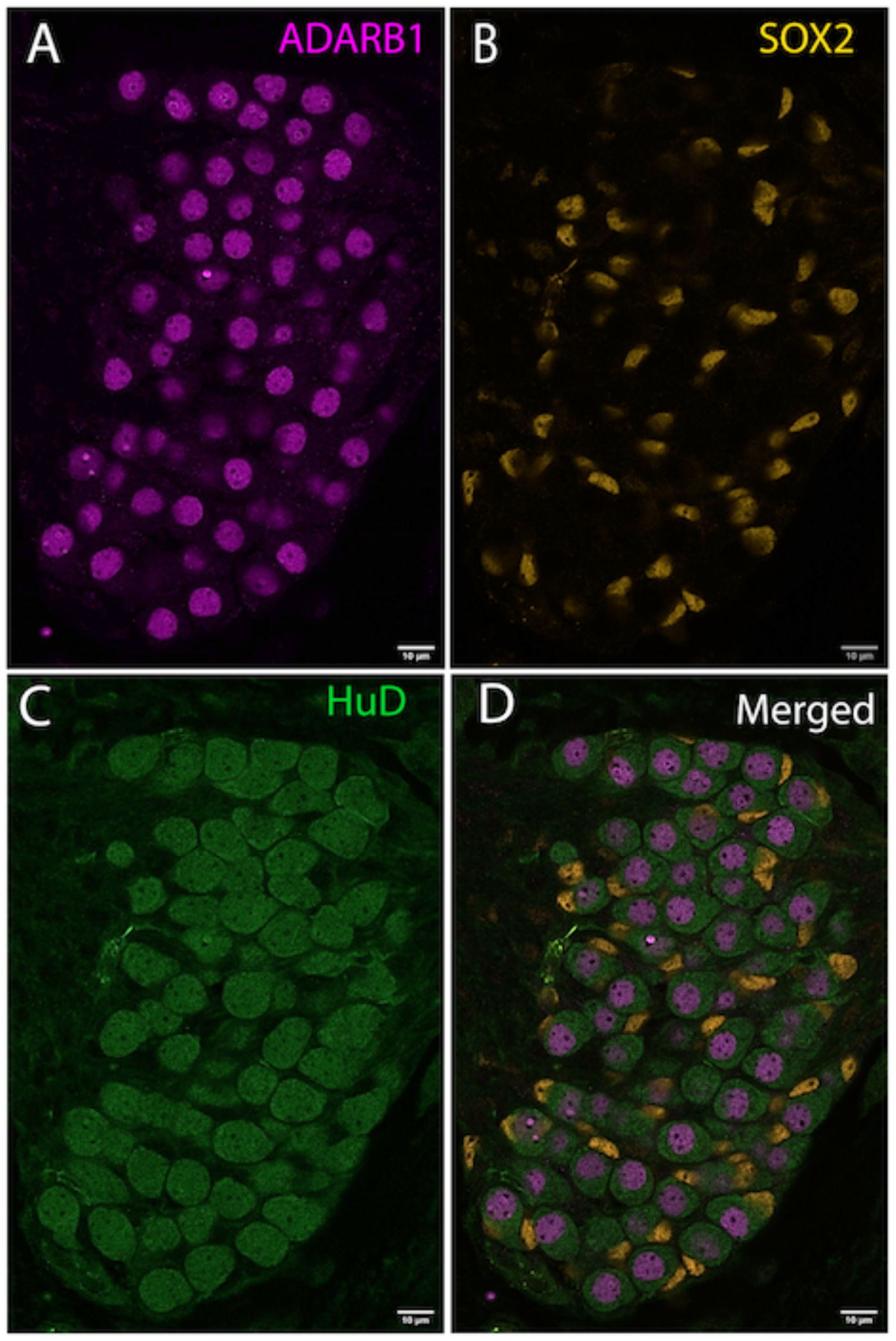
ADARB1 immunostaining is present in all HuD positive SGNs. (A-D) Confocal images of a SGN cross-section from a P21 mouse cochlea immunolabeled with ADARB1 (A, magenta), SOX2 (B, yellow) and HuD (C, green) clearly delineating the nuclear localization of ADARB1 as well as the nuclear and cytoplasmic distribution of HuD. The merged panel (D) Illustrates the colocalization of ADARB1-positive nuclei within HuD-positive SGNs and the lack of any colocalization with SOX2+ glial nuclei. Scale bar = 10 µm

**Figure 8:**
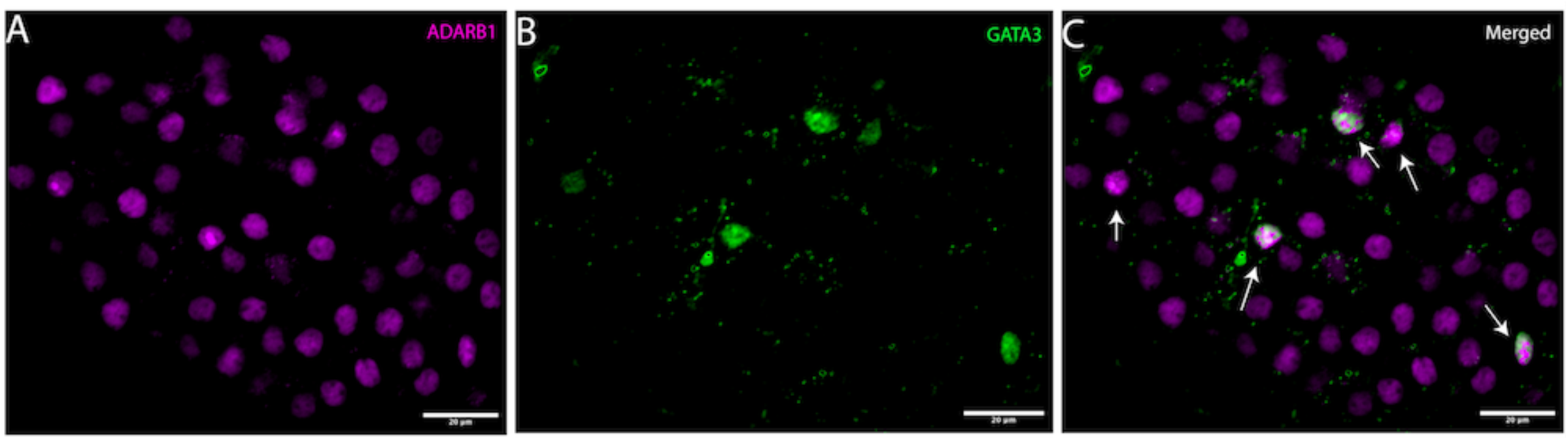
Colocalization of ADARB1-labeled nuclei with GATA3-labeled nuclei demonstrates that ADARB1 marks not only type I, but also type II, SGNs. (A-C) Confocal micrographs of a SGN cross-section from postnatal day 21 (P21) mouse cochlea immunolabeled with ADARB1 (magenta) and GATA3 (green) reveal colocalization of GATA3 in ADARB1-positive SGN nuclei (white arrows). Scale bar = 20 µm

### 4. Comparisons of manual NFH counts with automated ADARB1 counts demonstrate good agreement between the two methods

To assess the efficacy of ADARB1 as a marker for SGN quantification, we conducted a comparative analysis of manual NFH-based counts using traditional methods versus automated ADARB1 quantification in ImageJ as seen in Figure 9. This evaluation included 46 images obtained from temporal bone sections from six distinct CD1 mice all at age P21. Samples were immunolabeled for both ADARB1 and NF-H. Rosenthal’s canal was identified in each image and outlined and then separate images were created for each channel. A single, blinded investigator performed manual NFH-based neuronal counting, relying on subjective identification of stained neurons in the NFH-labeled images. A separate investigator quantified the positive nuclei in the ADARB1 images using the Process > binary > make binary and Analyze > analyze particles functions in ImageJ. Inter-rater reliability was assessed for the manually derived counts as compared with automated counts using Lin’s concordance correlation coefficient (CCC) and a Bland-Altman analysis (Figure 10). The Bland-Altman plot, executed in R with a range of +/-2SD, revealed a mean difference of 1.24 with an upper limit of +8.99 and a lower limit of −6.51. This suggests strong alignment between the two methods with ADARB1 quantification yielding marginally higher counts (by 1.24 per image on average) as compared to traditional manual NFH methods. Across the 41 images analyzed, the manual investigator recorded an average SGN count of 52.93, whereas the automated ADARB1 approach produced an average count of 54.17. Additionally, calculation of Lin’s coefficient of concordance demonstrated a strong agreement between the two quantification techniques, yielding a coefficient of 0.948. These results suggest that ADARB1 staining and automated SGN quantification are at least as reliable as conventional NFH-based manual counting and thus may provide a faster, more expedient approach without sacrificing accuracy.

**Figure 9:**
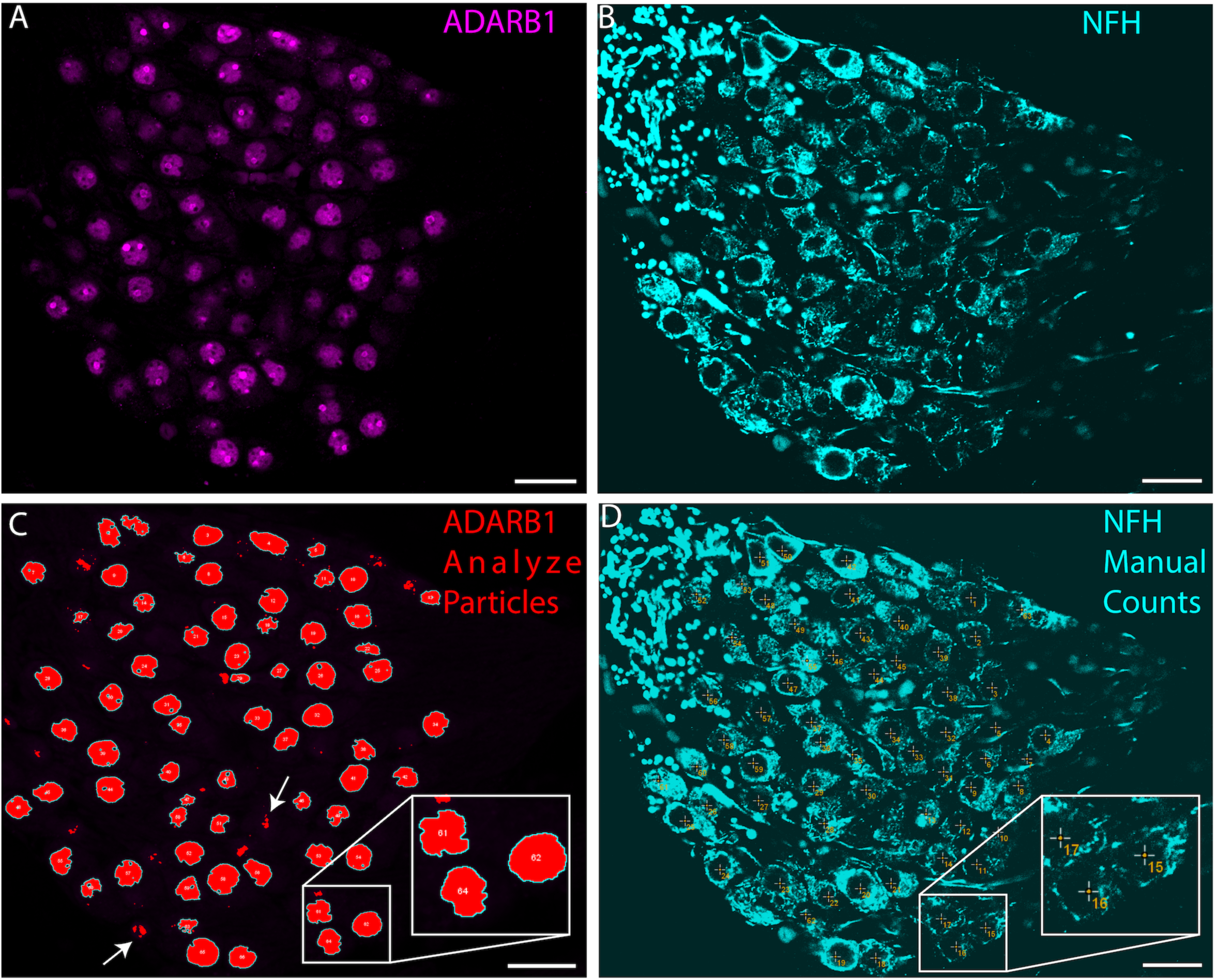
Comparison of automated ADARB1-based cell counts with manual NFH-based counts demonstrates the efficacy of ADARB1 for automated quantification of spiral ganglion neurons (SGNs). (A-D) Confocal micrographs of a cross-section from postnatal day 21 (P21) mouse cochlea immunolabeled with neurofilament heavy chain (NFH, cyan) and ADARB1 (magenta) are presented. (C) The result from an automated cell quantification utilizing ADARB1 immunolabeling, performed with the “Make binary” and “Analyze Particles” functions in ImageJ. Positively identified cells exhibit a cell border drawn in yellow. The white arrows show two areas not counted as they were excluded by size and circularity filtering in the Analyze Particles tool. (D) A conventional manual counting approach using NFH immunolabeling was undertaken by a blinded counter using the multi-point tool in ImageJ (numbered plus signs in RED), with a final count of NFH-63, ADARB1-66. Scale bar = 20 um

**Figure 10:**
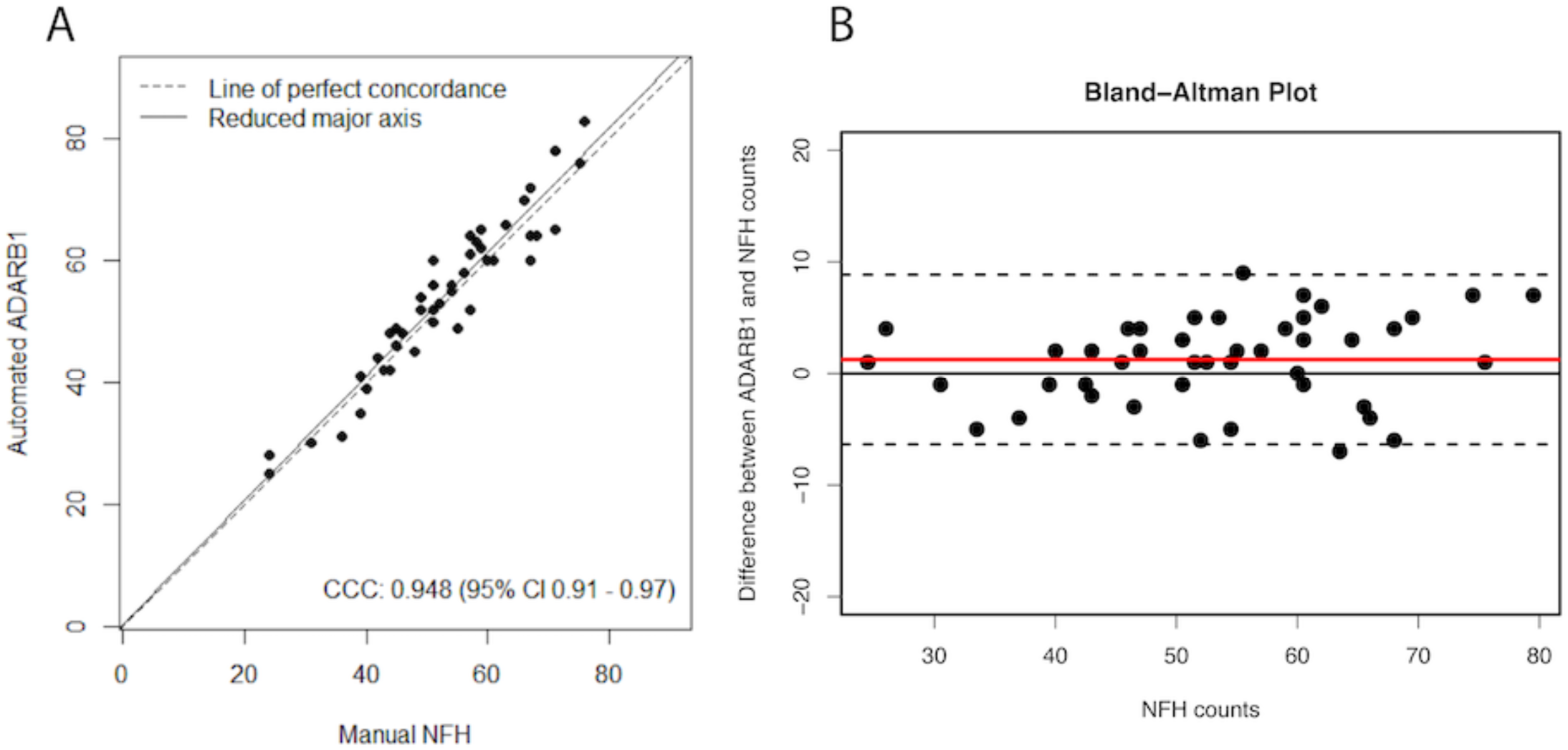
(A-B) Statistical analysis comparing manual counts using NFH and automated counts of ADARB1. A) Lin’s Concordance Correlation Coefficient (CCC) was calculated for 46 images of Rosenthal’s canal demonstrating a correlation of 0.948 and strong concordance between the two methods. B) A Bland-Altman plot was generated to help visualize concordance between the two methods. On average, the automated ADARB1 counts were higher than the manual NF-H counts by +1.24, which is shown as the bias line (red line). The upper limit (8.99) and lower limit (−6.51), shown as dashed lines, were calculated as two times the standard deviation around the mean (1.24) and suggest a high confidence level that future comparisons would lie within this range.

### 5. ADARB1 immunoreactivity in SGN nuclei persists to advanced ages

To assess the presence of ADARB1 in SGNs throughout the lifespan, we immunolabeled temporal bone sections from 2-year-old C57BL/6 mice. Across the samples investigated at this age, all of the NFH+ SGNs had robust ADARB1 immunolabeling in their nuclei (Figure 11), suggesting that ADARB1 immunofluorescence persists with aging. Of note, several images from higher frequency regions of the cochlea, where age-related hair cell loss was obvious, still exhibited robust ADARB1 labeling (Figure 12). These results suggest that ADARB1 may be valuable for SGN quantification throughout aging and in contexts where there is hair cell loss and reduced numbers of surviving SGNs (Figure 12).

**Figure 11:**
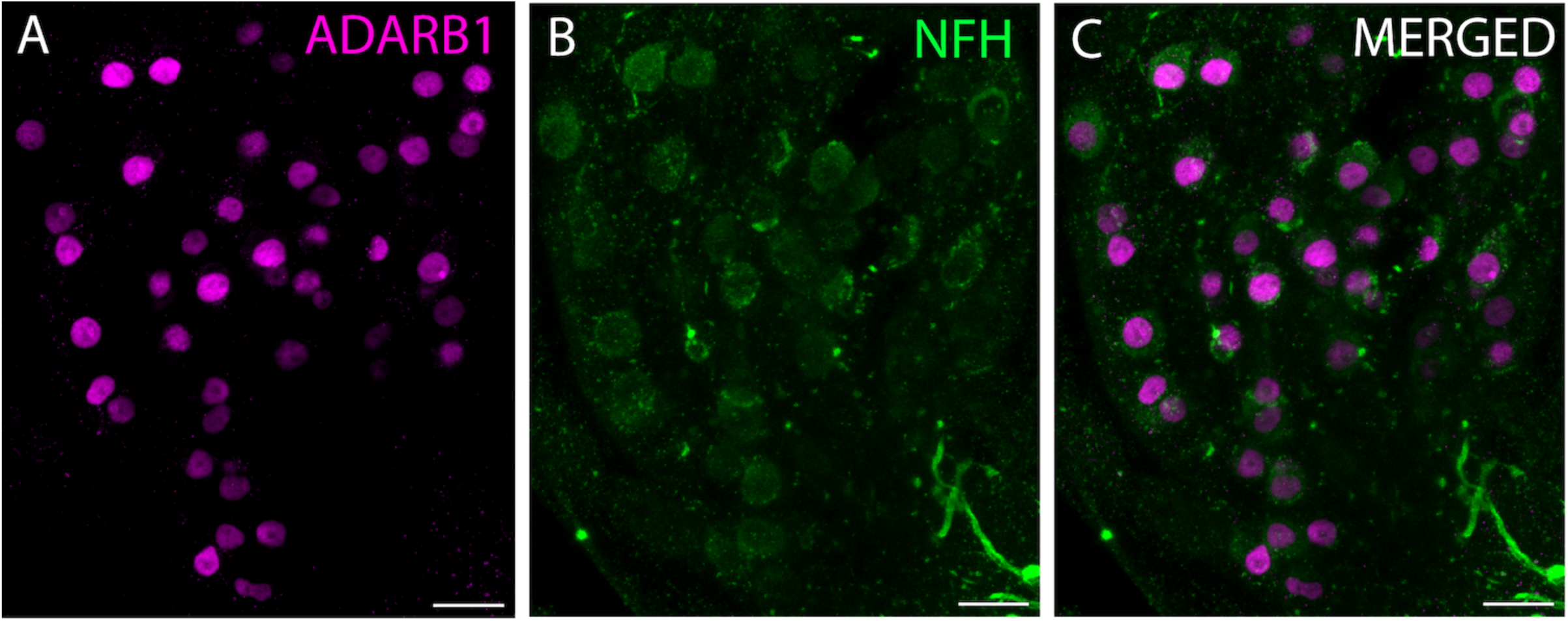
ADARB1 immunofluorescence persists in mouse SGNs throughout the lifespan. (A-C) ADARB1 and neurofilament heavy chain (NFH) immunolabeling in basal cochlear sections from 24-month-old mice show strong ADARB1 -immunoreactivity (magenta) in all NFH-positive SGNs (green). These findings further substantiate the reliability of ADARB1 as a marker for the quantitative assessment of SGNs in aged cochlear tissues..

**Figure 12:**
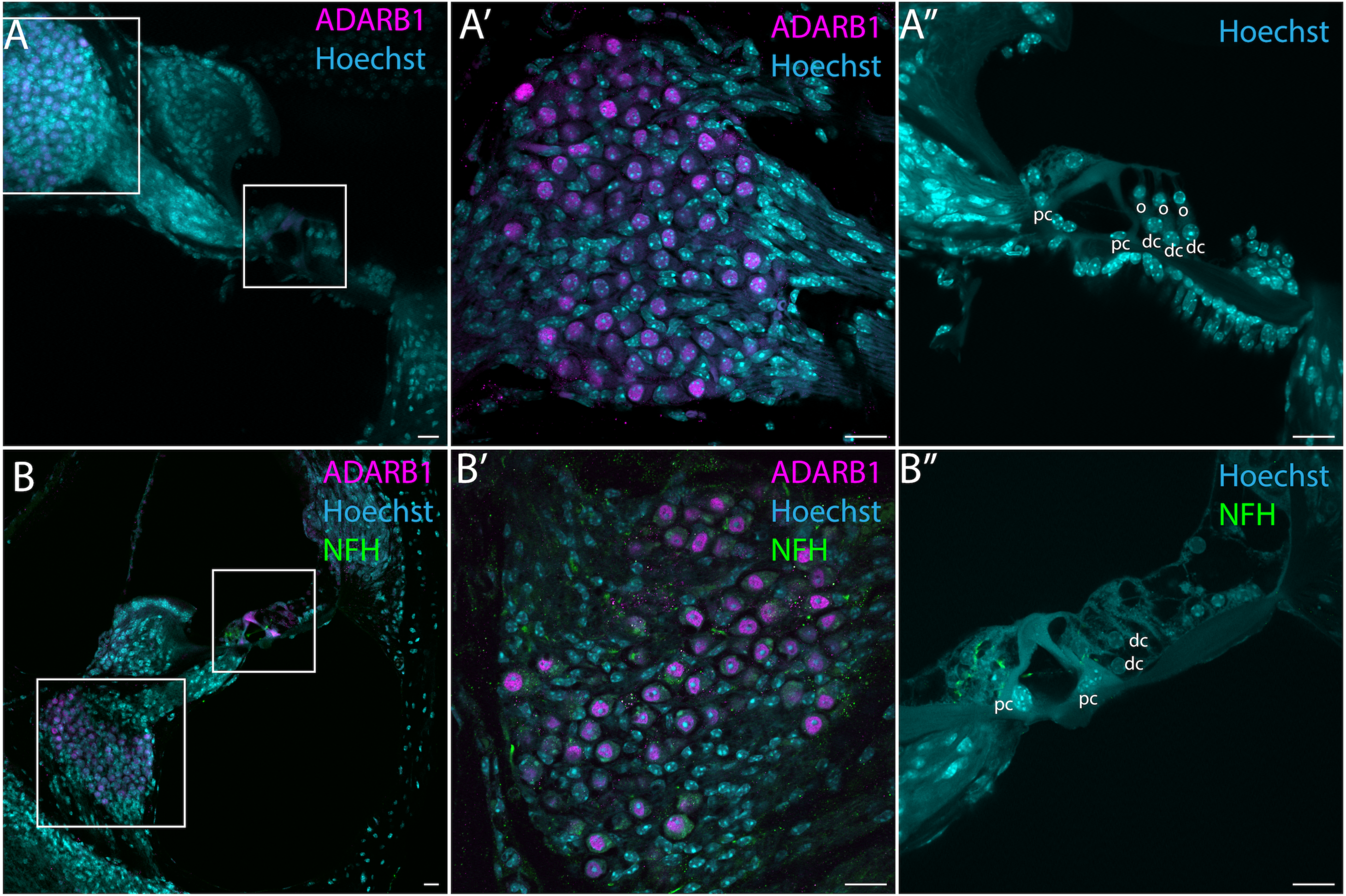
ADARB1 immunofluorescence is retained in surviving SGNs in aged (24-month-old) samples, despite hair cell loss. (A) A representative 20× image of the inner ear from a P21 mouse temporal bone is shown, with ADARB1 immunofluorescence (magenta) and Hoechst labeling (cyan). (A’) The SGN region from (A) is shown at 40X, as is the (A”) organ of Corti where the outer hair cells (o), Deiters cells(dc), and pillar cells (pc) can all be readily identified. (B) A 20X image from the mid-basal turn of a cochleae from a 24 month old C57Bl/6 mouse again showing ADARB1 (magenta) and Hoechst (cyan) labeling, as well as NFH (green). (B’) The SGN from (B) at 40X magnification shows robust ADARB1 labeling of the SGNs. (B”) A 40X image of the organ of Corti from (B) shows loss of outer hair cells characteristic of middle and basal turns from C57Bl/6 mice at this age. These findings support the use of ADARB1 as a reliable marker for SGNs in aged samples, even in circumstances where there is hair cell, and likely neuronal, loss. Scale bar 20 µm

### 6. ADARB1 is enriched in SGN nuclei in non-human primates and humans

To determine whether the findings reported above may be specific to mice, or if they may be more broadly applicable, we immunolabeled temporal bone sections from *Macaca fascicularis* and *Homo sapiens* to investigate ADARB1 immunoreactivity in these species. Our findings revealed robust SGN staining across these samples (Figures 13 & 14), though some variability was observed in the sections from human temporal bones. Specifically, while in many of the human samples, the overwhelming majority of SGN nuclei appeared to be labeled by the ADARB1 antibody (Figure 13), in some of the samples, not all of the cells had readily detectable ADARB1 immunofluorescence, and some exhibited nuclear immunofluorescence that was not dramatically brighter than the fluorescence of the surrounding soma (Figure 13). In some cases, the portion of SGNs that were clearly labeled for ADARB1 was only just above half of the total number of SGNs present in a given field of view. As noted in the methods, we were able to improve ADARB1 staining in many of these cases by increasing the concentrations of primary and secondary antibodies and by increasing incubation times as has been noted in other cadaveric temporal bone studies to improve immunolabeling (Lopez et al., 2016).Specifically, the primary antibody against ADARB1 (which worked well at concentrations of 1:200 in mice and macaques) showed some improvement of labeling in human sections when applied at 1:50 – 1:100. Similarly, increasing secondary antibody concentrations from 1:1000 to 1:500 improved signal in some samples, and increasing the duration of incubation in primary from overnight to ∼3 days and increasing the duration in secondary antibody from a few hours to overnight generally had positive effects as well. However, even under improved conditions, some of the samples still showed incomplete ADARB1 labeling where a majority of SGNs appeared labeled, but not always 100%. In contrast, ADARB1 immunostaining in the macaque samples was completely consistent with the pattern seen in mice which is to say that all of the SGN nuclei appeared to be clearly labeled across all of the samples examined (Figure 14). These results suggest that ADARB1 is readily expressed in human and macaque SGNs and that immunolabeling of ADARB1 in these species may be of use for automated SGN quantification. However, the results also suggest that sample quality is important. Unlike the mouse and macaque sections, the human temporal bone samples were of varying quality due to a number of factors such as heterogeneity in fixation, post-mortem time to fixation, storage conditions and duration, etc. These factors, which likely contributed to heterogeneous results in the human samples, are discussed in more detail below.

**Figure 13:**
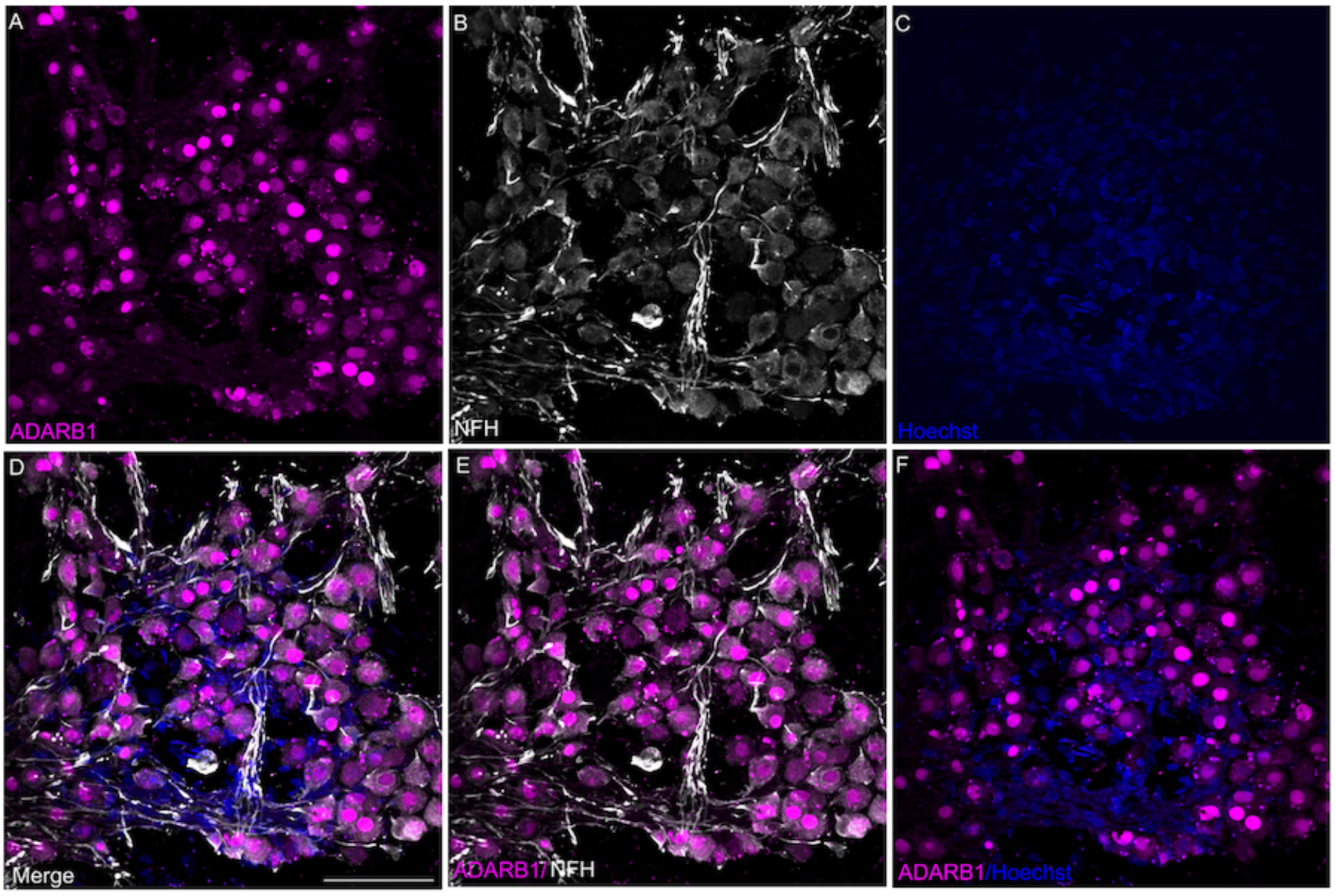
Immunofluorescence in cryosections from adult human temporal bones reveal ADARB1 is prominently expressed in the nuclei of SGNs. (A) A representative confocal image of adult human SGNs with strong nuclear labeling by anti-ADARB1 (magenta). (B)Neurofilamentous fibers labeling with anti-NF-H(white) and (C) Hoechst (blue) were used to identify the neurons and all cell nuclei, respectively. (D) Triple-labeling of ADARB1, NF-H, and Hoechst, as well as views of just ADARB1 and NF-H (E) and of just ADARB1 and Hoechst (F) highlight the localization of ADARB1 in the nuclei of SGNs. Images were taken at 40X magnification and scale bar = 500µm.

**Figure 14:**
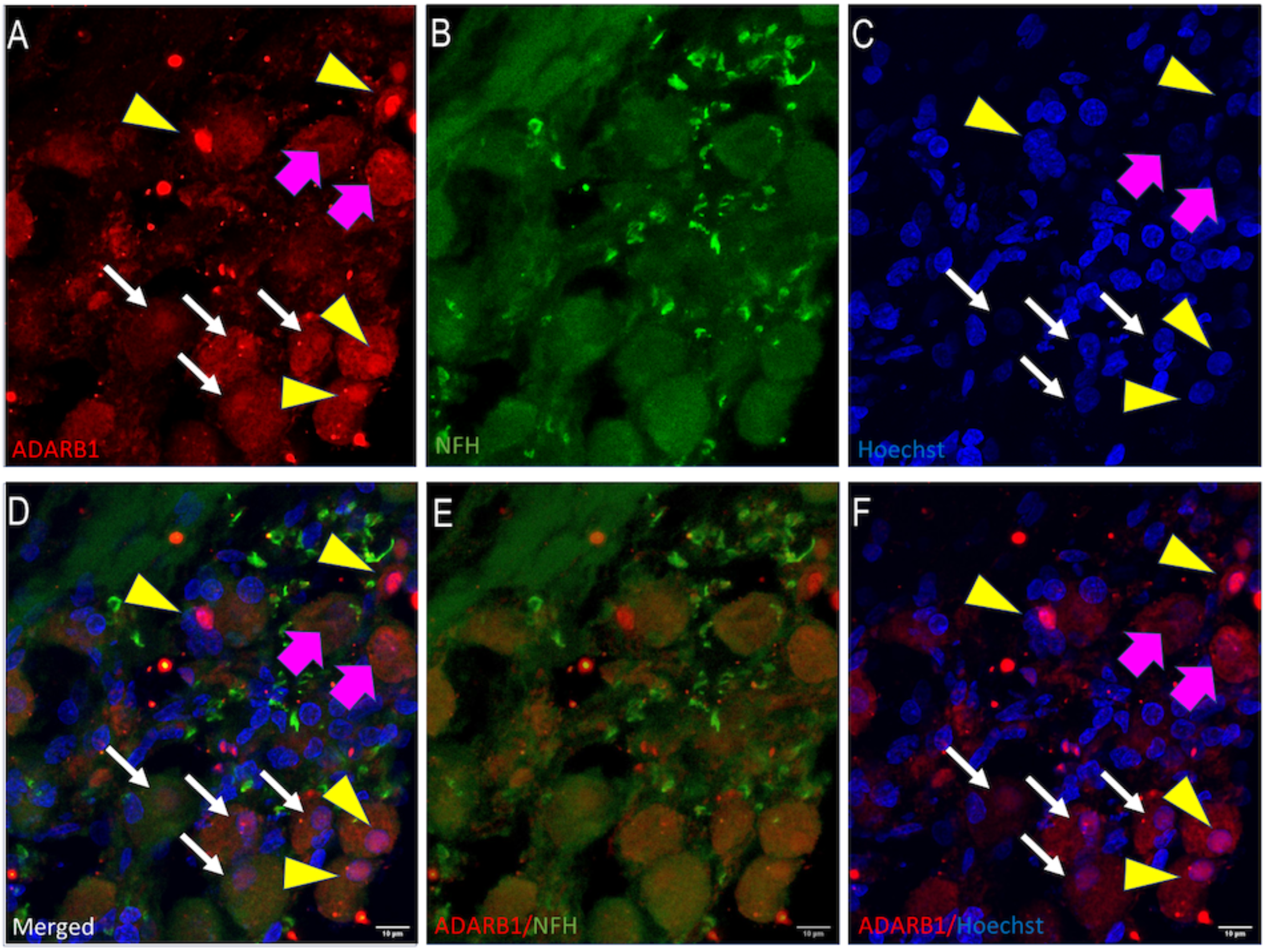
Some human cadaveric samples failed to exhibit consistent nuclear ADARB1 labeling in all of the SGN nuclei. (A) A confocal image from a less-well preserved human temporal bone section labeled with the antibody against ADARB1 (red). The presence of the ADARB1 can be observed in the SGNs, with several presenting clear enrichment in the cell nuclei (yellow arrowheads), some presenting only moderate enrichment in the nuclei (white arrows), and some appearing diffuse throughout the soma (magenta arrows). (B) NFH labeling (green) was used to confirm neuronal identify. (C) Hoechst (Blue) was used to label the cell nuclei. (D) In an image where the Hoechst, NFH, and ADARB1 signals are merged, SGN nuclei with less prominent ADARB1 label can be discerned (white arrows). (E) Double-labeling images of ADARB1 with NFH or of ADARB1 with Hoechst (F) provide additional views. Images were taken at 40X magnification and scale bar = 10um

**Figure 15:**
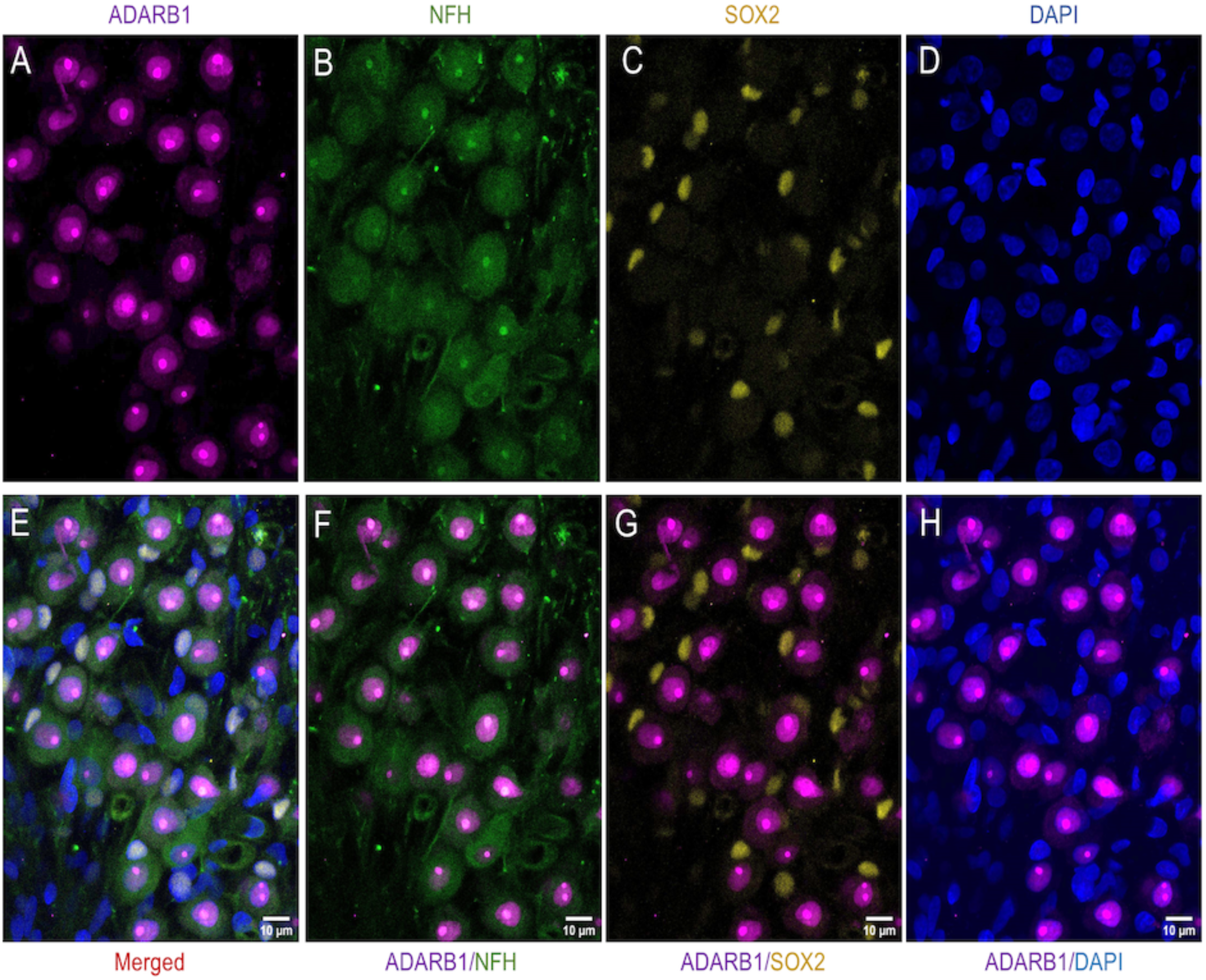
A confocal image of a macaque temporal bone section labelled with anti-ADARB1(A, Magenta), anti-NFH (B, Green), anti-SOX2 (C, yellow), and Hoechst (D, Blue). A merged image with all of the channels (E), and subsequent panels showing two-channel overlapping views, clearly show that ADARB1 immunoreactivity is selectively within NFH-positive SGNs (F), is not prominently expressed in SOX2-positive glial nuclei (G), but is nuclear (H) within the SGNs. Scale bar = 10um

## Discussion

### ADARB1 expression in the spiral ganglion: potential roles for synaptic transmission, development, function, and dysfunction

We hypothesized that ADARB1 (aka ADAR2) would be expressed in spiral ganglion neurons due to the fact that the afferent synapses express the GluA2 receptor subunit which needs to be edited in order to render the receptors calcium impermeable. For vertebrates, glutamate serves as the principal excitatory neurotransmitter, mediating the activation of afferent neurons such as spiral ganglion neurons (SGNs) within the inner ear. Transmission of auditory information from sensory hair cells of the organ of Corti to SGNs is facilitated via the activation of AMPA receptors. These receptors are tetrameric complexes composed of subunits GLUA1–4. Notably, the GLUA2 subunit is of particular interest, as its incorporation into the AMPA receptor complex modulates calcium permeability, with normally edited GLUA2 limiting calcium permeability (Burnashev et al., 1992; Sommer, Köhler, et al., 1991; Sommer, Kohler, et al., 1991). It has been shown previously that GLUA2 is edited by ADARB1 and that, in the absence of this editing, AMPA receptors remain calcium permeable leading to seizures and perinatal lethality (Higuchi et al., 2000). Reports of RNA seq data from the inner ear suggest that *Adarb1* RNA is highly expressed in SGNs and here we have confirmed that pattern for both the *Adarb1* transcript and the ADARB1 protein using RNAscope probes and an ADARB1-specific antibody that has been previously validated by knockout (Soundararajan et al., 2022a), knockdown(Wang et al., 2019), and overexpression(Soundararajan et al., 2022a). We further demonstrate that ADARB1 immunolabeling is enriched in the SGN cell nuclei and that this pattern of labeling lends itself to automated SGN quantification, as demonstrated here with the free, open-access software ImageJ (or Fiji). While not the main focus of the work presented here, it is of potential importance that ADARB1 is highly expressed in the SGNs and appears to be upregulated as the neurons mature and acquire auditory responsiveness. This pattern of increasing expression as neurons mature is consistent with what has been previously published for excitatory glutamatergic neurons in the central nervous system (Matho et al., 2021), and is also consistent with RNA-seq data from SGNs in mice which suggests increasing expression of *Adarb1* developmentally, with plateaued expression from P8 onward (Li et al., 2020). As expression of *Gria2*, which codes for the GluA2 subunit, appears to be already high in SGNs at E15.5 with only modest increases in expression thereafter (Li et al., 2020), this suggests that developing, immature SGNs may present with some meaningful proportion of unedited, calcium permeable (CP-) GluA2 receptors. As calcium permeable AMPA receptors have been implicated in synaptic plasticity, development, and various disease states (Cull-Candy & Farrant, 2021; de León-López et al., 2025; Henley & Wilkinson, 2016; Man, 2011), the results here suggest hypotheses regarding SGN development and function which could represent novel areas of investigation for the field of hearing research. For example, the editing of CP-GluA2 subunits to impermeable GluA2 subunits has been shown to be important for the high level of mapping that occurs in the visual system(Liu et al., 2019)and an analogous function in the auditory system would imply GluA2 editing as potentially important for the establishment of tonotopy in the developing cochlea. Furthermore, GluA2 editing has been implicated in neurodevelopmental and neurodegenerative conditions (Konen et al., 2020; Wright & Vissel, 2012)suggesting intriguing hypotheses about the potential role for GluA2 editing in injury or disease states of the inner ear such as after noise or blast trauma, iatrogenic damage accompanying cochlear implants, Alzheimer’s Disease, and possibly other conditions (Salpietro et al., 2019; Wright et al., 2023). For example, GluA2 Q/R editing is modulated by changes in ADAR2 levels that occur during ischemia (Peng et al., 2006) and in response to excitotoxic levels of glutamate (Mahajan et al., 2011). Furthermore, evidence indicates that reducing calcium permeability by inhibiting non-GluA2 AMPA receptors confers protection against noise-induced synaptopathy in SGNs(Hu et al., 2020) This finding suggests a novel mechanistic role for calcium modulation, which could be affected by inhibition of RNA editing as a potential target in the investigation of otoprotective strategies. Finally, while GluA2 is the most well-known target of ADARB1 editing, it is possible, even likely, that other transcripts in SGNs are edited by this enzyme. For instance, ADAR2 also edits GluK2 (Gurung et al., 2018), and transcripts necessary for neurite growth and guidance have recently been shown to undergo A-to-I editing (La Via et al., 2025) suggesting other developmental and functional aspects of SGNs may be influenced by ADARB1 expression and activity. Thus, our confirmation of the presence of high levels of ADARB1 in the SGNs has the potential to open several new avenues of study in the inner ear

### The utility of ADARB1 immunolabeling for SGN quantification

Manual counting of spiral ganglion neurons (SGNs) has been widely used in hearing research to study aging, neuropathies, traumatic brain injury (TBI), and other pathologies, but it is historically a time-consuming and labor-intensive process. We investigated ADARB1 as a potential alternative SGN marker, hypothesizing that its nuclear-specific staining would improve neuron segmentation and facilitate automated cell counting. Unlike conventional markers that label the neurites and/or soma, and contribute to crowded views of SGNs and overlapping signals in micrographs, ADARB1 staining enhances resolution, enabling more effective discrimination of neurons, and easier automated or manual counting. Our comparison of manual counting using NFH and automated counting with ADARB1 showed highly similar results, supporting ADARB1 as a reliable and efficient alternative for SGN quantification. The adoption of this nuclear marker offers several advantages, primarily in terms of labor and time savings, though possible additional benefits include reduced requirements for expertise and training, potential limiting of investigator bias, as well as greater flexibility in experimental design, image acquisition, and analysis. Here, we used a traditional method involving cochlear sectioning and analysis with the simple, freely available Analyze Particles function in ImageJ, enabling faster, more consistent analysis than manual approaches. However, with a robust and reliable nuclear marker such as ADARB1, tissue clearing and lightsheet microscopy become a more tangible prospect. Similarly, large z-stacks of images can be acquired using light-sheet, confocal, or structured-illumination approaches, and automated SGN quantification can then be undertaken using other software options for 3D analysis, such as Napari, IMARIS, Amira-Avizo, or others.

ADARB1 labeling to identify SGNs can also reduce the need for expertise in inner ear anatomy, allowing new students or other novices to accurately count cell numbers. However, success in this regard depends on good sample preparation and staining, as well as image quality. Thus, expertise is likely still required for dissection, sectioning, staining, and image acquisition. Indeed, some of the samples used in this study still showed low but variable background or non-specific fluorescence, damage to sections from dissection or handling, spurious signals from foreign matter (e.g., dust or precipitates), and some edge artifacts. In instances where image quality was suboptimal, we found that a semi-automated rather than fully automated approach was helpful. Preprocessing steps, available via ImageJ tools or plug-ins, such as background subtraction, binary erosion, and watershed processes, could be applied to reduce background fluorescence or signal bleed-through and help with segmentation when nuclei at different depths within tissue samples are still in close proximity or present as overlapping in maximum intensity projections. In cases with spurious signals or edge artifacts, the polygon outline tool was helpful for clearing signals located outside Rosenthal’s canal, and adjusting the particle size range in the analyze particles function helped to remove noise. In our dataset, the lower limits of particle size ranged from 9 µm² to 14 µm², depending on image quality and background interference.

ADARB1 immunoreactivity in juvenile (P21) mice could represent an idealized case for SGN quantification. As such, we tested ADARB1 labeling in other contexts, including in aged mice, archived Macaque temporal bone samples, and archived human cadaveric samples (also generally from aged individuals). ADARB1 immunoreactivity not only persists in SGN nuclei of temporal bones from 24-month-old mice, but was robust within the basal cochlear turn, including in samples with obvious hair cell loss. While these regions in the basal turns of aged mouse cochleae also exhibited some SGN loss, strong ADARB1 nuclear labeling was observed in all NF-H-positive SGNs, suggesting that ADARB1 is reliable for detecting all surviving SGNs. This result suggests that ADARB1 immunostaining is useful for studies on cochlear aging and for those involving hair cell loss or neurodegeneration. Though of note, it was often helpful to increase imaging parameters, such as laser power, brightness, and contrast, to achieve ADARB1 fluorescence levels in 24-month-old mice that were comparable to those readily detected in P21 mice. This requirement suggests a potential decrease in ADARB1 signal intensity in aged SGNs, which may reflect reduced expression, though any such changes in expression across age would require future experiments to validate. Additionally, ADARB1 shows nuclear localization in SGNs from both non-human primate and human specimens, highlighting its potential translational relevance. Although variability was observed among human samples, likely attributable to differences in fixation protocols and tissue storage, signal strength remained sufficient to support its feasibility as a marker in these tissues. Further studies are warranted to optimize and validate ADARB1-based quantification across species and pathological states. However, as we note below, there are likely to be scenarios where ADARB1 labeling may be less than ideal for SGN quantification including in tissue from the developing inner ear and in samples such as cadaveric sections where fixation and storage conditions cannot be tightly controlled. Still, being able to automate or semi-automate SGN counting with ADARB1 labeling should significantly reduce the time and resources spent by many researchers currently relying on manual quantification.

### Level of evidence and limitations of the study

Although our data strongly suggest the utility of ADARB1 immunolabeling for the automation of counting of spiral ganglion neurons, there are some limitations. As with all immunolabeling experiments, the quality of the samples, the consistency of the procedures, the validity of the antibody, and the resulting image acquisition are critical factors. When immunolabeling, there is always the possibility of background, incomplete filtration of emitted light (a.k.a. bleed-through), and non-specific signal which can be caused by spurious binding, edge artifacts, or particulates such as dust or precipitates in antibody solutions. Also, heterogeneity of fluorescence intensity both within and across nuclei can lead to variability when thresholding images to yield binary data for analysis. This can make it difficult for software such as the ImageJ Analyze Particles function to accurately count neurons without some level of manual involvement or pre-processing. Here we found the noise removal and erode functions (under the Process tab) in ImageJ to be helpful in generating processed images that worked well with minimal changing of settings within the Analyze particles function. However, modification of settings such as minimum particle size and circularity within the Analyze particles interface could also be used to achieve the desired filtering. With images of high quality, these steps were unnecessary, but for some images, these more semi-automated approaches were helpful. We also observed variable quality of ADARB1 labeling and detection in the human samples which we suspect was likely due to a lack of control over the fixation process and the storage of tissue in the weeks or months immediately post-mortem. As the tissue had been sourced from cadavers used in medical education, embalming methods and quality were variable and all cadavers were maintained in anatomy tables at room temperature for at least two months while in use for medical education prior to temporal bone removal, decalcification, and sectioning. In some of the human samples, perhaps only 50-60% of the SGNs were labeled by the ADARB1 antibody. Increasing the concentration and duration of primary antibody application generally improved the staining, but there were still some samples where even though ADARB1 appeared to label the majority of SGNs, some ADARB1 negative SGNs still persisted. We and others have previously noted similar difficulties with the immunostaining of cadaveric samples, even when using other more established markers (Ghosh et al., 2022; Lopez et al., 2016). While it is also possible that time in storage (5+ years in −80°C) might influence ADARB1 immunoreactivity, the macaque samples used in this study were stored identically and for the same amount of time as the human samples, and in all of the macaque sections 100% of the SGNs were labeled by the ADARB1 antibody suggesting that the time in storage was not likely to be the primary issue. In fact, the robust ADARB1 immunofluorescence still readily obtainable in sections archived for so long was a surprising, but positive finding. Still, we cannot, on the current evidence, rule out biological causes for the observed heterogeneity in the human samples such as possible variability of expression in human ears due to age or other factors. However, the much more consistent results in the mice and macaques suggest that tissue fixation, sample preparation, and room temperature storage are likely to be factors contributing to reduced efficiency of immunolabeling in cadaveric sections and we hope that future studies with better preserved human specimens will be able to validate this hypothesis.

Another potential limitation with regard to the use of ADARB1 immunolabeling for SGN quantification is the variation in ADARB1 expression with age, particularly prior to hearing onset. Although we are cautiously optimistic that ADARB1 could still be a useful tool for counting SGNs in the first few postnatal weeks in mice, we suspect it may be more challenging during development or perinatally. Indeed, as noted above, RNAseq data suggest *Adarb1* expression reaches its maximum in SGNs at P8 in mice suggesting that anti-ADARB1 immunolabeling could be highly specific to SGNs at that time. However, this remains unconfirmed as we did not examine any tissue from mice between the ages of P0 and P21. Furthermore, we did not investigate any samples from macaques or humans at any embryonic or developmental timepoints, thus we can only speculate that ADARB1 labeling would be less robust at these stages based on extrapolation from the mouse data. With regard to advanced age, we discovered that robust ADARB1 immunofluorescence persists well into adulthood, particularly in mice and macaques, but likely in human SGNs as well. However, we used two different mouse strains for the 24-month and young adult groups without direct, within-strain comparisons across ages, and, as noted, we did not achieve 100% labeling of SGNs with anti-ADARB1 across all of our human samples which mostly came from aged individuals. Thus, the potential exists for diminished labeling with extreme aging which may limit utility in some contexts (e.g., human cadavers), but may also be important for investigating RNA editing in SGNs during aging or in other neurodegenerative conditions. Future experiments will be needed to fully validate ADARB1 labeling and/or expression in these contexts.

With regard to the antibodies used in this study, all of the antibodies used for co-labeling with ADARB1 (i.e., SOX2, HuD, NFH, GATA3, NeuN), have been previously published and validated to label the cell types as reported here (Berglund & Ryugo, 1991; Coate & Kelley, 2013; Vanevski & Xu, 2015). For ADARB1 labeling, we initially tested two different antibodies, but proceeded with the more extensive characterization outlined here using only one of these. The antibody that is reported here (Proteintech 22248-1-AP) was chosen as it yielded more robust labeling of SGN nuclei at a lower concentrations than the alternative (Sigma #SAB1405426), though both antibodies labeled SGN nuclei in temporal bones from 1–2-month-old mice. Furthermore, the documentation for both antibodies include validation by Western blotting and either ICC or IF with the mouse antibody having also been validated with transfected and non-transfected cells and the rabbit antibody being validated via immunoprecipitation experiments. Furthermore, the antibody used primarily in this report has been documented in at least 16 publications including multiple reports where *Adarb1* was knocked out, knocked down, and/or overexpressed with the corresponding manipulations being measurable via immunoblotting with the antibody (Kokot et al., 2022; Soundararajan et al., 2022b). When considered in addition to our ability to localize *Adarb1* transcripts to SGNs via RNAscope in P21 mice, the level of evidence for anti-ADARB1 in SGN quantification is fairly high, particularly with high quality samples where fixation and storage conditions are well-controlled.

## Conclusions and future directions

Robust, nuclear localized ADARB1 immunofluorescence provides for high confidence of efficient SGN counting with reduced crowding and the potential for automation. Additionally, independent labeling with Sox-2 confirmed that ADARB1 does not colocalize with glial cells, while co-labeling with HuD, NFH, and GATA3 indicated its presence in both type I and type II neurons. Furthermore, existing RNAseq data and our own RNAscope data (Figure 2), when coupled with the immunolabeling, strongly suggest *Adarb1* is highly expressed in SGNs at both young and old adult ages, raising intriguing questions for further exploration in auditory research. Future studies could investigate what other transcripts are edited in SGNs, as well as how GluA2 calcium permeability changes during development, in response to insults, or with aging.

## Notes

### Competing Interest Statement

The authors have declared no competing interest.

